# Serum metabolomic biomarkers of perceptual speed in cognitively normal and mildly impaired subjects with fasting state stratification

**DOI:** 10.1101/2020.09.03.282343

**Authors:** Kamil Borkowski, Ameer Y. Taha, Theresa L. Pedersen, Philip L. De Jager, David A. Bennett, Rima Kaddurah-Daouk, John W. Newman

## Abstract

Cognitive decline is associated with both normal aging and early pathologies leading to dementia. Here we used quantitative profiling of metabolites involved in the regulation of inflammation, vascular function, neuronal function and energy metabolism, including oxylipins, endocannabinoids, bile acids, and steroid hormones to identify metabolic biomarkers of mild cognitive impairment (MCI). Serum samples (n =210) were obtained from subjects with or without MCI opportunistically collected with incomplete fasting state information. To maximize power and stratify the analysis of metabolite associations with MCI by the fasting state, we developed an algorithm to predict subject fasting state when unknown (n =71). In non-fasted subjects, linoleic acid and palmitoleoyl ethanolamide levels were positively associated with perceptual speed. In fasted subjects, soluble epoxide hydrolase activity and tauro-alpha-muricholic acid levels were negatively associated with perceptual speed. Other cognitive domains showed associations with bile acid metabolism, but only in the non-fasted state. Importantly, this study shows unique associations between serum metabolites and cognitive function in the fasted and non-fasted states and provides a fasting state prediction algorithm based on measurable metabolites.

## 1. Introduction

Neurocognitive disorders including Alzheimer’s dementia (AD) are associated with cognitive decline. Biochemical markers of altered cognitive capacity may provide diagnostic and prognostic biomarkers of these diseases and their associated metabolic trajectories before clinical symptoms manifest. Additionally, such biomarkers could provide new insights into the mechanisms of cognitive decline. Cognition can be decomposed into dissociable domains, characterized as perceptual speed, perceptual orientation along with semantic, working and episodic memory. These cognitive domains become increasingly inter-correlated as people become cognitively impaired ^1^, and have been linked to pathologic changes in the brain ^2^. While the events which initiate these changes are as yet unknown, dysregulated cellular mechanisms associated with metabolic dysfunctions and/or inflammatory responses are attractive hypotheses.

It has recently become clear that cardiometabolic disorders and associated low-grade systemic inflammation and altered lipid and energy metabolism, are risk factors for AD ^3–5^. Therefore, changes in circulating markers of low-grade inflammation and metabolism may track these pertinent metabolic changes. Obesity and the metabolic syndrome shift the profile of both plasma lipids and multiple lipid-derived physiological mediators ^6,7^. Four important families of such lipid mediators readily detected in the circulation are the oxygenated polyunsaturated fatty acids (i.e. oxylipins), the endogenous cannabinoid receptor activators and their structural equivalents (i.e. endocannabinoids), bile acids and steroids.

The oxylipins including fatty acid alcohols, diols, epoxides, ketones, and prostanoids are derived from multiple polyunsaturated fatty acids (PUFA) by the action of cyclooxygenases (COX), lipoxygenases (LOX), cytochrome P450 (CYP), soluble epoxide hydrolase (sEH) or reactive oxygen species (ROS) and various downstream enzymatic processes ^8,9^. Circulating endocannabinoids are produced either by acylation and release of acyl ethanolamides from phosphatidylethanolamine, or as a product of glycerol-lipid metabolism (monoacylglycerols).

Oxylipins and endocannabinoids are known to regulate multiple processes including both acute and low-grade systemic inflammation ^9,10^, cardiovascular health ^11^, neuronal outgrowth, cell differentiation and energetics ^12^. Bile acids and steroid are also linked to the regulation of glucose and insulin metabolism ^13^, energy metabolism and inflammation ^14,15^ and implicated in the pathogenesis of type 2 diabetes and metabolic syndrome ^16^. Previous studies reported associations between AD, cognition and plasma levels of oxylipins ^17^, bile acids ^18,19^ and steroids ^20,21^. However, broader simultaneous assessments of lipid mediator profiles in the context of mild cognitive impairment have not been conducted to date.

Frozen collections of serum and plasma from studies of neurocognitive disorders, including measures of cognitive function, provide a resource for biomarker discovery in this arena ^22^ However, opportunistically collected samples rarely contain information regarding fed/fasted states, which can compromise “omics” analyses. Here, we took advantage of data and biospecimens from subjects in the Religious Order Study and Rush Memory and Aging Project (ROS/MAP) ^23^, develop a predictive tool for the fasted/non-fasted state discrimination and stratify our biomarker discovery effort by the fasted state. We describe an exploration of circulating oxylipin, endocannabinoids, bile acids, and steroids for biomarkers of cognitive impairment, providing insights into unique associations in basal and postprandial metabolism.

## 2. Materials and methods

### 2.1 Subjects

Participants in the Religious Orders Study (ROS) are older nuns, priests, and brothers from across the United States, while those in the Rush Memory and Aging Project (MAP) are older lay persons from the greater Chicago area ^23^. Both studies enrolled persons without known dementia and perform annual detailed clinical evaluations. Both studies were approved by an Institutional Review Board of Rush University Medical Center. All participants signed an informed consent and a repository consent to allow their biospecimens and data to be shared. ROS/MAP resources can be requested at www.radc.rush.edu. The current sample consists of 196 subjects with 14 subjects having two blood samples collected on average 5.8 ± 3.3 years apart. Repeated blood draws were in opposite fasting states (either fasted or non-fasted). Subjects demographics: 22% male, 95% white and non-Hispanic. Average age (mean ± standard deviation) = 78.2 ±7.2, average BMI = 27.2 ±4.8 average years of education = 15.3 ±2.8. Number of known fasted samples as recorded by a technician = 59; non fasted = 80, unknown = 71.

### 2.2 Clinical evaluation of cognition

All subjects are under a yearly structured clinical evaluation, including a medical history, neurologic examination and cognitive testing. The studies have 19 tests in common. Eleven tests are used to inform diagnostic classification of dementia and its causes, and cognitive impairment with as previously reported ^24,25^. Mild cognitive impairment (MCI) refers to people with cognitive impairment without dementia ^24^. No cognitive impairment (NCI) are those without dementia or MCI ^24^. Seventeen tests are used for four measure of global cognition and five distinct cognitive domains including perceptual speed, perceptual orientation, episodic memory, semantic memory and working memory. The global cognition was calculated by converting each test to a z score based on the baseline mean and standard deviation and averaging the 17 tests; the domains were created by averaging subsets of z-scores as previously reported in detail^26^.

### 2.4 Quantification of clinical lipids, glucose and glycosylated hemoglobin

Phlebotomists and nurses collected the blood specimen as previously reported ^27^. Tests were performed by Quest Diagnostics (Secaucus, NJ). For this study we used glucose (mg/dL), hemoglobin A1c, expressed as a percentage of hemoglobin, and a basic lipid panel consisting of total cholesterol, HDL and LDL cholesterol, and triglycerides (all units mg/dL).

### 2.5 Quantification of oxylipins, endocannabinoids, PUFA, non-steroidal anti-inflammatory drugs, bile acids and steroids

Serum concentrations of non-esterified PUFA, oxylipins, endocannabinoids, a group of non-steroidal anti-inflammatory drugs (NSAIDs) including ibuprofen, naproxen, acetaminophen, a suite of conjugated and unconjugated bile acids, and a series of glucocorticoids, progestins and testosterone were quantified by liquid chromatography tandem mass spectrometry (LC-MS/MS) after protein precipitation in the presence of deuterated metabolite analogs (i.e. analytical surrogates) using modifications of published procedures ^28,29^. Samples were processed with rigorous quality control measures including the analysis of batch blanks and replicates of serum pools and NIST Standard Reference Material 1950 (Sigma-Aldrich). Samples were re-randomized for acquisition, with method blanks and internal reference material and calibration sets scattered regularly throughout the set. Instrument limits of detection (LODs) and limits of quantification (LOQs) were estimated according to the Environmental Protection Agency method (40 CFR, Appendix B to Part 136 revision 1.11, U.S. and EPA 821-R-16-006 Revision 2). These values were then transformed into sample nM concentrations by multiplying the calculated concentration by the final sample volume (i.e. 250 μL) and dividing by the volume of sample extracted (i.e. 50 μL). Using the Students t-Distribution, the t-value was determined at a 95% 1-tail confidence level to define the LOD. A complete analyte list with their LOD and LOQ is provided in the **Supplemental Table S1.**The majority of analytes were quantified against analytical standards with the exception of eicosapentaenoyl ethanolamide (EPEA), palmitoleoyl ethanolamide (POEA), and the measured PUFA [i.e. linoleic acid (LA); alpha-linolenic acid (aLA); arachidonic acid (AA); eicosapentaenoic acid (EPA); docosahexaenoic acid (DHA)]. For those compounds the area counts were recorded, adjusted for deuterated-surrogate and the relative response factors were expressed as the relative abundance across all analyzed samples. MAGs are reported as the sum of 1 and 2 isomers, due to their potential isomerization during the sample processing. The complete metabolomic data are available via the AD Knowledge Portal (https://adknowledgeportal.synapse.org). The AD Knowledge Portal is a platform for accessing data, analyses, and tools generated by the Accelerating Medicines Partnership (AMP-AD) Target Discovery Program and other National Institute on Aging (NIA)-supported programs to enable open-science practices and accelerate translational learning. The data, analyses and tools are shared early in the research cycle without a publication embargo on secondary use. Data is available for general research use according to the following requirements for data access and data attribution (https://adknowledgeportal.synapse.org/DataAccess/Instructions). See doi: 10.7303/syn22344904.

### 2.6 Statistical analysis

All statistical tests were performed using JMP Pro 14 (JMP, SAS institute, Carry, NC). Prior to analysis, two data points were removed as outliers using the robust Huber M test and missing data were imputed using multivariate normal imputation for variables which were at least 75% complete. Imputation facilitated multivariate data analysis and did not significantly influence univariate results. Additionally, variables were normalized, centered and scaled using Johnson’s transformation, with normality verification using the Shapiro-Wilk test. Cognitive sores were adjusted for BMI, sex, age, race and education and their residuals were used for further analysis. Metabolite inter-correlations were evaluated using Spearman’s rank-order correlations. Variable clustering by hierarchical cluster analysis used the Ward agglomeration. Multiple comparison control was accomplished with the false discovery rate (FDR) correction method of Benjamini and Hochberg 30, with the number of independent observations determined by the correlative structure of variables (number of variable clusters).

Predictive models for fasting state and cognitive functions were prepared using a combination of bootstrap forest and stepwise linear regression modeling, with Bayesian information criterion (BIC) cutoff. Variables most frequently appearing in the models were identified by bootstrap forest (logistic or regression, respectively): trees in forest = 100; terms sampled per split = 5; bootstrap sample rate = 1. A variable contribution scree plot was generated using variable rank and the likelihood ratio of chi-square (for categorical fasted/non-fasted prediction) or sum of squares (for continues cognitive scores). The scree plot was used to determine a likelihood ratio of chi-square or sum of squares cutoff value for variables contributing to the model. Selected variables were then subjected to stepwise logistic regressions for fasted/non-fasted predictions, or stepwise linear regressions for cognitive scores. Data were split into training (60%) and validation (40%) cohorts, with balanced separation across metabolites and cognitive domains. Stepwise analysis was performed with the maximal validation r^2^; as the model stopping criteria, or if an additional step increased the BIC.

## 3. Results

### 3.1. Serum lipid mediators predict the fasting state

Our cohort consists of 210 samples including 59 fasted, 80 non-fasted and 71 of unknown fasting state. Using samples with known fasting state, A fasting state prediction model was developed using measured PUFA, lipid mediator, bile acid, steroid, clinical lipid and glucose data. Clinical lipids (e.g. triglycerides or cholesterol) and glucose did not produce strong predictive models and did not contribute to the final model. A high probability of the fasted state was described by low levels of the LA-derived CYP metabolite [12(13)-EpOME], low levels of the primary conjugated bile acid glycochenodeoxycholic acid (GCDCA) and elevated levels of the glycine-conjugated oleic acid (NO-Gly; **Fig. 1 A and B**). The model misclassification rate was 12%., with fasting probability described by the **Equation 1** and **Equation 2**.

**Figure 1:**
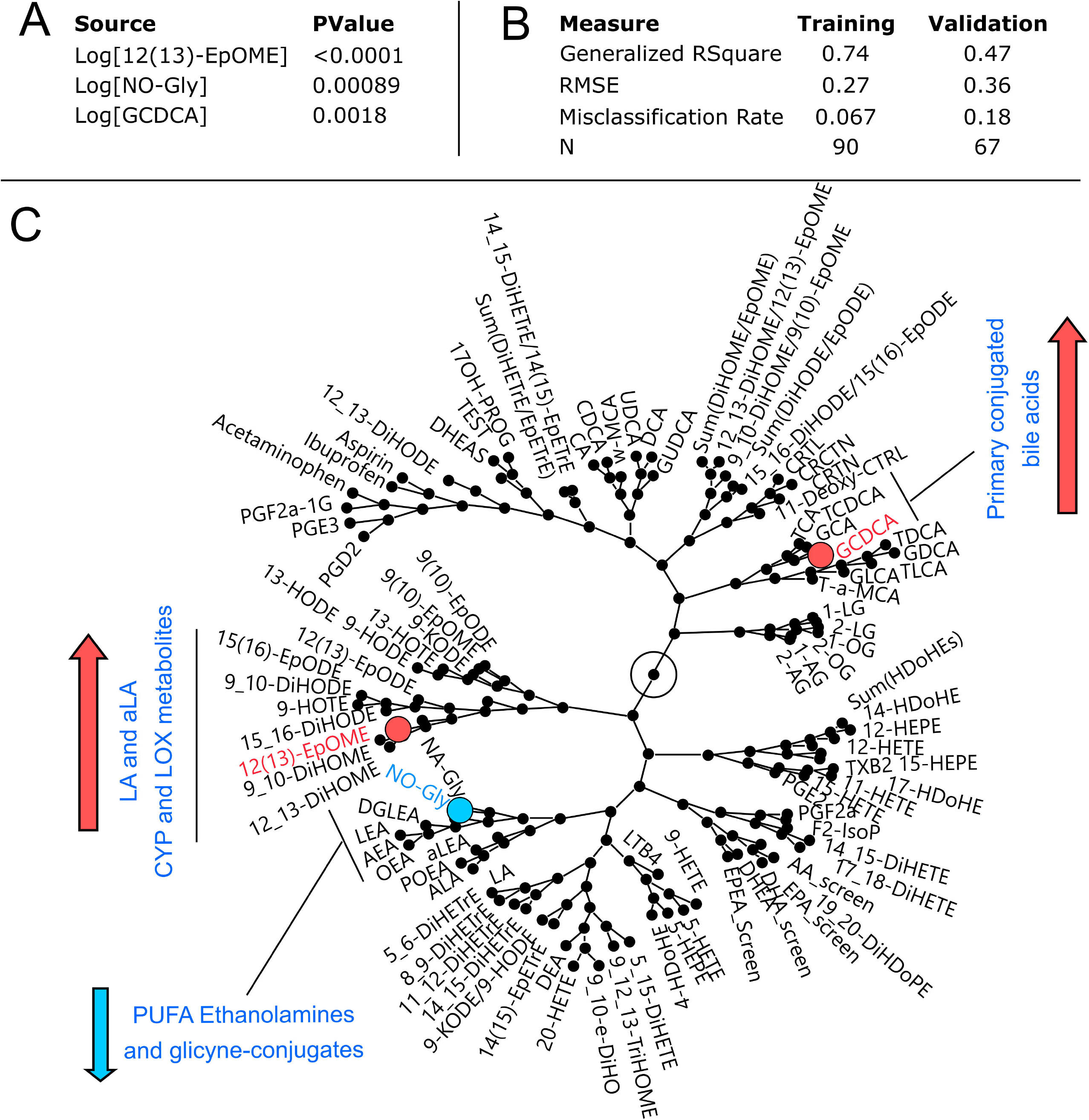
Serum lipid metabolites and bile acids are predictors of the fasting state. **A)**Stepwise logistic model parameters predicting the fasting state using 12(13)-EpOME, GCDCA and NO-Gly. **B)**Visualization of the correlative environment (generated using hierarchical clustering) of metabolites used for fasting state prediction. Nodes represent branching points in the hierarchical clustering network with metabolites on the fringe named. Metabolite used in the final model are indicated by colors. Directionality of changes in metabolites due to non-fasted state compared to the fasted state are indicated by arrows.

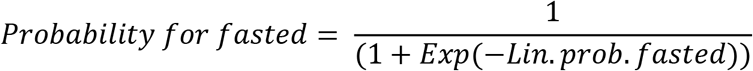

**Equation 1**. Probability of the fated state. Where “Lin.prob.fasted” is defined by the Equation 2:

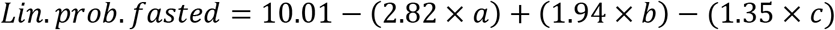

**Equation 2**. Lin prob fasted: *a* = Log[12(13)-EpOME]; *b* = Log(NO-Gly); *c* = Log(GCDCA). Concentrations expressed in (nM).

Oxylipins, endocannabinoids, PUFA, bile acids and steroids create correlative structures along metabolic pathways or from common precursor fatty acids (**Fig. 1C**). Therefore, similar fasting state predictions could be achieved by substituting metabolites with ones close in the correlation network. For example, NO-Gly can be effectively replaced by oleoyl ethanolamide (OEA). Validation of model was performed using an independent cohort ^31^ of fasted plasma (n =133) and showed a misclassification rate of 17%, dropping to 12% when considering samples with a probability of prediction >70%. To facilitate understanding of oxylipin and endocannabinoid metabolic relationship, their synthesis pathway from PUFA as well as coverage of metabolites detected in this study are presented in the **Supplemental Fig. S1**.

### 3.2. Fasted and non-fasted serum reveal distinct associations between lipid mediators and cognitive functions

Spearman’s rank correlations demonstrated associations between serum lipid mediators and cognitive functions. Cognitive scores were adjusted for BMI, gender, age, race and education. The analysis was stratified by subject fasting states. **Figure 2** shows correlation between the five cognitive domains.

**Figure 2.**
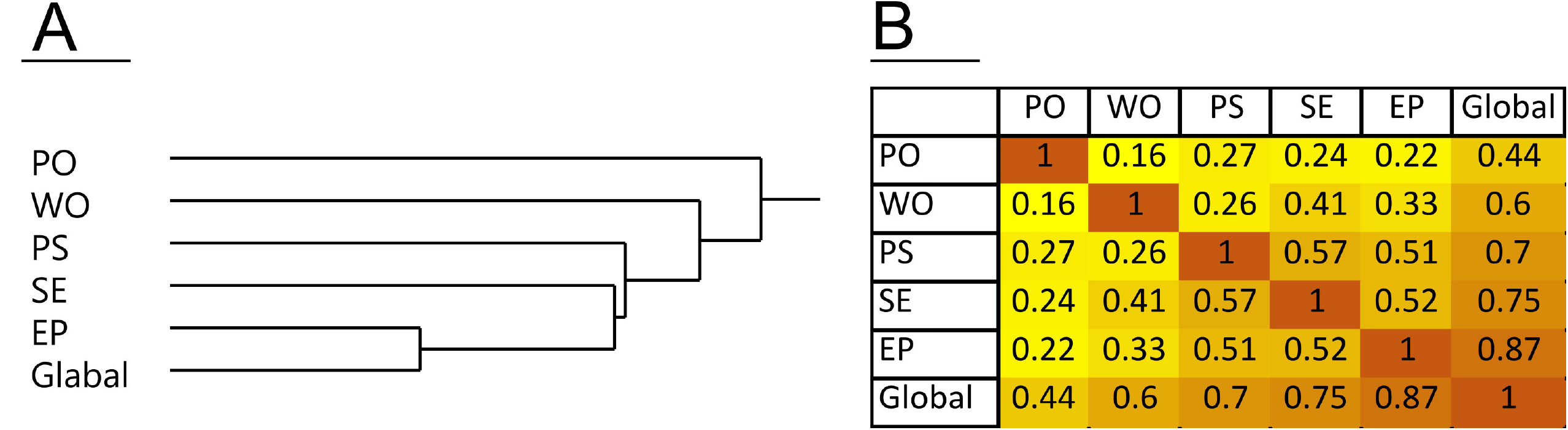
Correlative relationships between cognitive domains. A) Hierarchical clustering of cognitive domains using Ward method. B) Pearson’s correlation matrix. PO – perceptual orientation; WO – working memory; PS – perceptual speed; SE – semantic memory; EP – episodic memory; Global – global cognition.

Oxylipins and endocannabinoids showed the greatest number of associations with perceptual speed (from 8% to 10% of metabolites in fasted and non-fasted samples respectively, **Table 1**). The number of associations for other cognitive domains and global cognition did not exceed 5% of the measured oxylipins and endocannabinoids (**Supplemental Table S2**). Fasted and non-fasted samples showed distinct correlation patterns. In non-fasted subjects perceptual speed was positively associated with the level of free PUFA, particularly LA, eicosapentaenoic acid (EPA) and docosahexaenoic acid (DHA), as well as the N-acyl ethanolamides derived from palmitoleate (POEA), and EPA (EPEA) and the EPA- and DHA-derived mono-alcohols (15-HEPE and 4-HDoHE respectively). These associations were absent in fasted subjects. Additionally, when fasted and non-fasted subjects were analyzed together without fasting state stratification, the above-mentioned associations were either not present or weaker than in non-fasted subjects alone, see **Supplemental Table S3**).

**Table 1.**
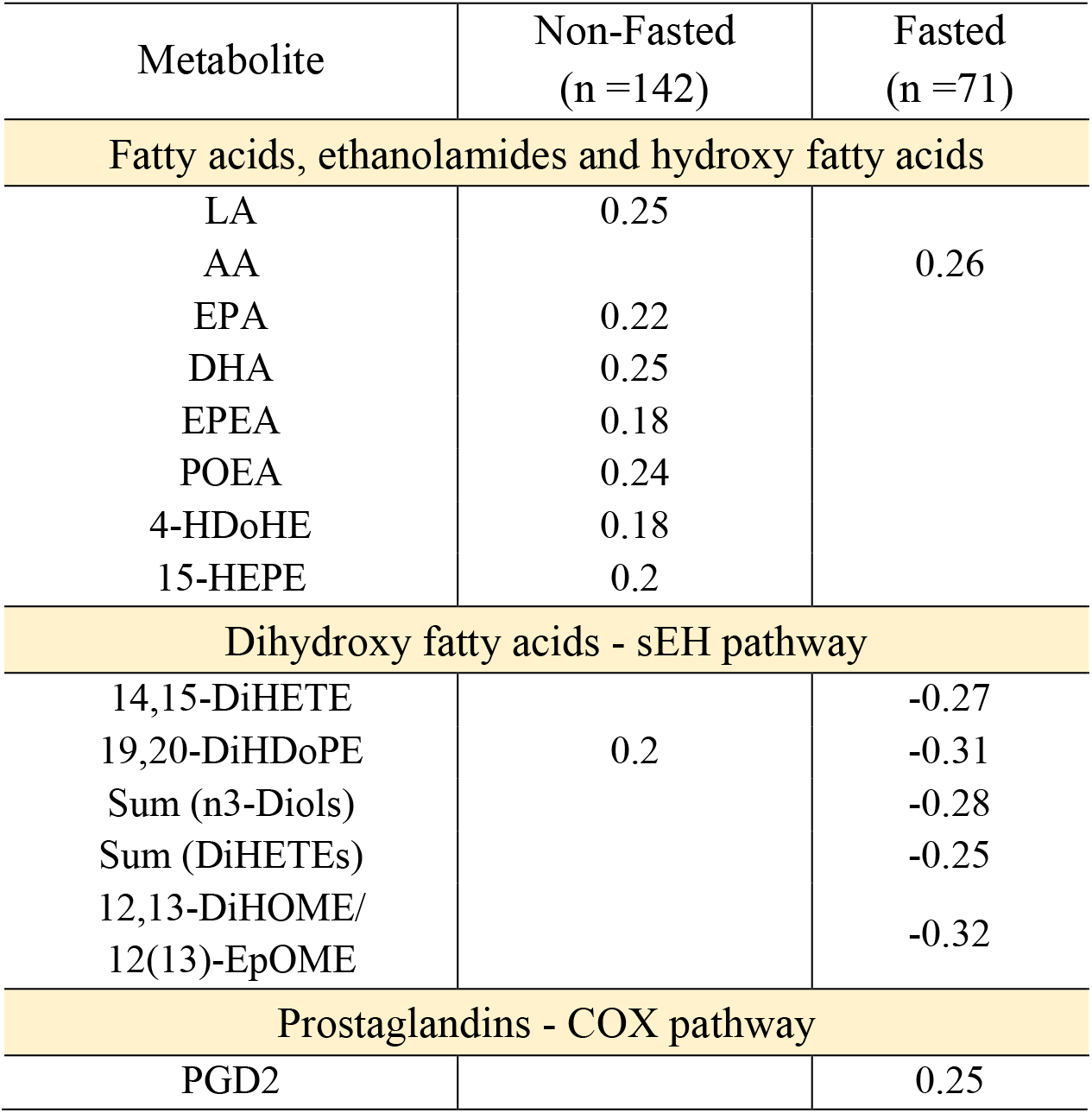
Spearman’s rank order correlations between serum oxylipins and endocannabinoids and perceptual speed. The numbers represent Spearman’s ρ with the p value <0.05 and FDR corrected with the q =0.2. Full names of all identified compounds are presented in the **Supplemental Table S1** and correlation for all cognitive domains are presented in the **Supplemental Table S2**.

On the other hand, fasted samples manifested negative correlations between perceptual speed and sEH products of EPA and DHA, and the ratio of LA vicinal diols (i.e. those with two hydroxy groups on adjacent carbons) to their corresponding epoxides, an estimator of sEH activity ^32^. This association was not detected in non-fasted subjects. Importantly, the cognitive domains scores were not different between the fasting states. Additionally, interaction with sex were not detected for the above-mentioned associations.

Numerous significant correlations were detected between bile acid levels and cognitive scores, mainly in non-fasted subjects (episodic memory: 9% to 38%; semantic memory: 3% to 25%; global cognition: 6% to 25%; and perceptual speed: 3% to 16% in fasted and non-fasted subjects respectively, **Table 2**). Perceptual orientation and working memory showed <6% associations (**Supplemental Table S2**).

**Table 2.** **Spearman’s rank order correlations between serum bile acids and cognitive domains**. The numbers represent Spearman’s ρ with the p value <0.05 and FDR corrected with the q =0.2. Full names of all identified compounds are presented in the **Supplemental Table S1** and correlation for all cognitive domains are presented in the **Supplemental Table S2**. PS – perceptual speed; SE – semantic memory; EP – episodic memory; Global – global cognition.

| Metabolite | Non-Fasted (n=142) |  |  |  | Fasted (n=71) |  |  |  |
| --- | --- | --- | --- | --- | --- | --- | --- | --- |
| Cognitive domain | PS | SE | EP | Global | PS | SE | EP | Global |
| Bile Acids - unconjugated |  |  |  |  |  |  |  |  |
| CDCA | 0.2 | 0.19 |  |  |  |  | 0.27 |  |
| DCA |  | 0.2 |  |  |  |  |  |  |
| Bile Acids - conjugated |  |  |  |  |  |  |  |  |
| TCDCA |  | -0.2 |  |  |  |  |  |  |
| TLCA | -0.2 | -0.27 | -0.28 | -0.29 |  |  |  |  |
| TDCA |  |  | -0.18 |  |  |  |  |  |
| GDCA |  |  | -0.21 |  |  |  |  |  |
| Bile acids - conjugated/unconjugated |  |  |  |  |  |  |  |  |
| TDCA/DCA |  | -0.25 | -0.18 | -0.22 |  |  |  |  |
| GDCA/DCA |  | -0.3 | -0.23 | -0.27 |  |  |  |  |
| GCDCA/CDCA | -0.24 | -0.28 |  | -0.2 |  |  |  |  |
| GCA/CA |  |  | -0.22 |  |  |  |  |  |
| TCA/CA |  |  | -0.21 |  |  |  |  |  |
| GUDCA/UDCA |  |  |  |  |  |  | 0.3 | 0.32 |
| Bile acids - glycine/taurine |  |  |  |  |  |  |  |  |
| (GDCA+GLCA)/<br>(TDCA+TLCA) |  |  | 0.18 | 0.19 |  |  |  |  |
| Bile Acids - dehydroxylation by bacteria |  |  |  |  |  |  |  |  |
| TDCA/CA |  |  | -0.26 | -0.19 |  |  |  |  |
| GDCA/CA |  |  | -0.28 | -0.18 |  |  |  |  |
| DCA/CA |  |  | -0.19 |  |  |  |  |  |
| GLCA/CDCA | -0.19 |  |  |  |  |  |  |  |
| TLCA/CDCA | -0.24 | -0.28 | -0.24 | -0.27 |  |  |  |  |
| Bile Acids - other |  |  |  |  |  |  |  |  |
| T-a-MCA |  |  |  |  | -0.28 | -0.26 | -0.29 | -0.31 |
| T-a-MCA/CDCA | -0.22 | -0.27 |  | -0.2 |  | -0.26 | -0.35 | -0.31 |
| w-MCA/T-a-MCA |  | 0.23 |  | 0.2 |  | 0.28 |  |  |

In non-fasted subjects, unconjugated bile acids correlated positively with perceptual speed and semantic memory. On the on the other hand, conjugated bile acids and the ratios of conjugated to unconjugated bile acids showed negative associations with perceptual speed, semantic and episodic memory and global cognition. Additionally, positive associations were observed between the ratio of glycine to taurine conjugated bile acids and episodic memory and global cognition. Negative associations were observed between the ratio of the downstream product to their precursor - cholic acid (CA) and episodic memory and global cognition. Finally, negative associations were observed between the ratio of tauro-alpha-muricholic acid (T-a-MCA) and its precursor chenodeoxycholic acid (CDCA).

Few associations between cognition and bile acids were observed in the fasted subjects. Negative associations were observed between T-a-MCA and T-a-MCA/CDCA ratio and episodic and semantic memory, perceptual speed and the global cognition. Also, positive associations were observed between the ratio of glycine conjugated to unconjugated ursodeoxycholic acid (UDCA) and episodic memory and global cognition. No associations were found between cognitive domains and steroid hormones.

### 3.3. Fasted state lipid mediators predict perceptual speed

Predictive models revealed covariate relationships between serum lipid mediators and cognition. Stepwise linear regression models (**Supplemental Table S4)**were built independently for each cognitive domain and for fasted/non-fasted samples. Valid models could not be generated using non-fasted subject data. Consistent with Spearman’s correlation results, perceptual speed formed the strongest model (R^2^_perceptual speed_ =0.44; R^2^_perceptual orientation_ =0.4; R^2^_episodic memory_ = 0.29; R^2^_global cognition_ =0.24) using samples from fasted subjects. The final model for perceptual speed is presented in the **Fig. 3.** This model included the ratio of LA-derived 12,13-DiHOME to 12(13)-EpOME, the sum of n-3 diols, consisting of EPA- and DHA-derived diols (14,15-DiHETE, 17,18-DiHETE and 19,20-DiHDoPE) and T-a-MCA. The epoxide/diol ratio and the sum of n-3 diols contributed the most to the model, with p-values of 0.0012 and 0.0007 respectively, and T-a-MCA with a weaker, but significant contribution (p value = 0.046). **Supplemental Fig. S2** shows correlative structure of all detected metabolites in fasted subjects. Sum of n-3 diols consist of all detected EPA and DHA diols. Corresponding EPA and DHA-derived epoxides were not detected.

**Figure 3:**
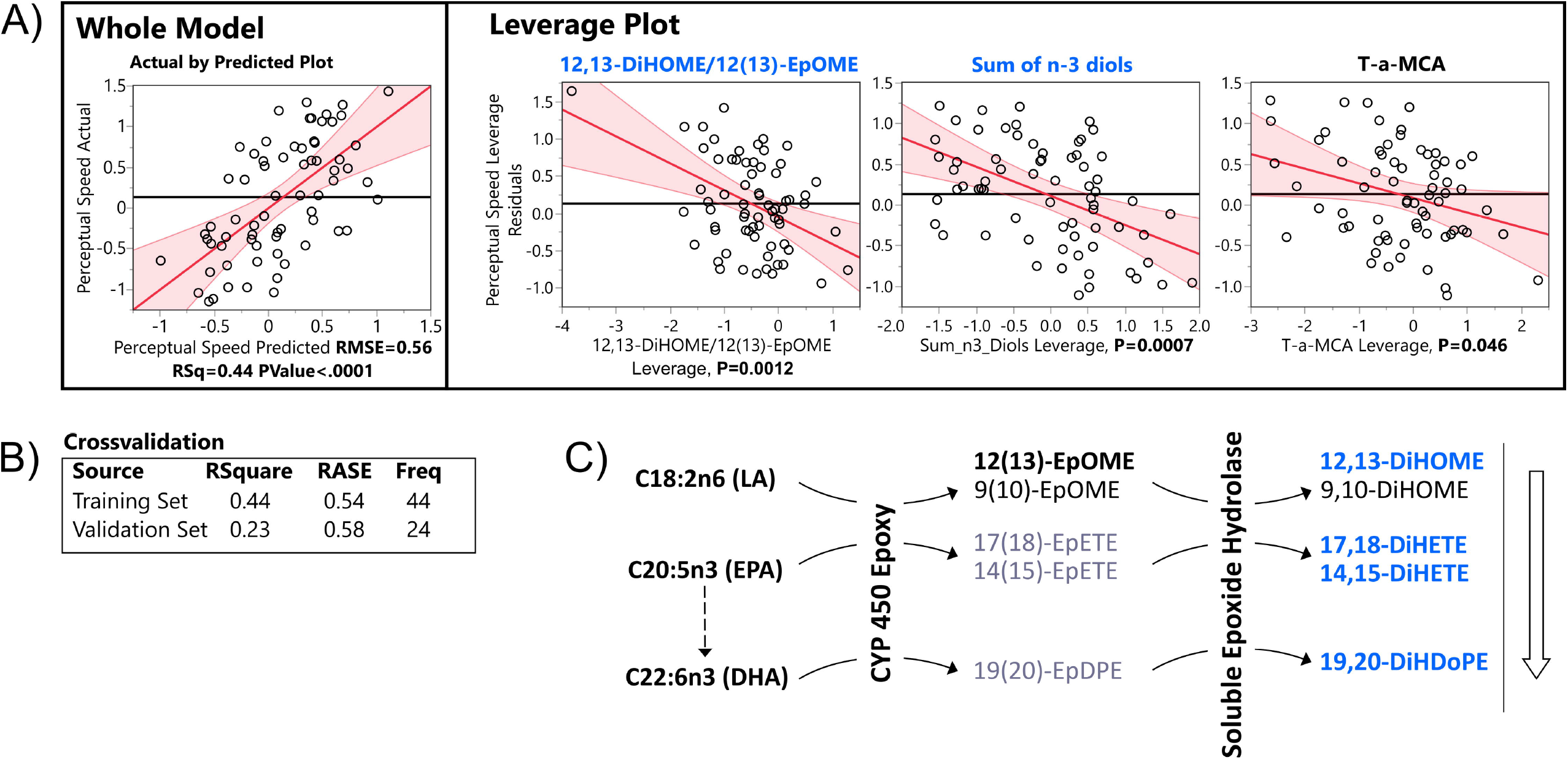
Least square regression model of perceptual speed. **A)** Actual by predicted plot of a whole model and leverage plots of model components. **B)** Model cross-validation statistics using training set (60%, n =44) and validation set (40%, n =33). **C)** Model components of soluble epoxide hydrolase metabolism projected onto their metabolic pathway. Metabolic pathway starts with the fatty acids on the left, farther, metabolizing enzymes are indicated on the arrows. Multiple possible metabolites of the pathway are indicated. Metabolites of sEH used for the model are highlighted. Color of the metabolites as well as an arrow next to the metabolic pathway represents directionality of the correlation with perceptual speed (orange – positive, blue – negative). RMSE – root mean squared error, LA – linoleic acid, CYP 450 – cytochrome p450, sEH – soluble epoxide hydrolase, EpOME – epoxy octadecanoic acid, DiHOME – dihydroxy octadecanoic acid, EpETE – epoxy eicosatrienoic acid, DiHETE – dihydroxy eicosatrienoic acid, EpDPE – epoxy docosapentaenoic acid, DiHDoPE – dihydroxy docosapentaenoic acid.

## 4. Discussion

In the current study we identified serum lipid mediators associated with cognitive function in a cohort exhibiting normal to mildly-impaired cognition. Moreover, this study provides a solution to the unknown fasting state of subjects that may occur when using opportunistically collected samples, and identifies unique associations with cognition in both fasted and non-fasted states.

Opportunistically collected serum and plasma are often collected without regards an individuals’ fasting state, compromising investigations probing peripheral factors influenced by postprandial fluctuations in the metabolome ^33^, proteome ^34^ and transcriptome ^35^. Using metabolomic data, we have developed a tool to determine subject fasting states and show enhanced statistical power with fasting state stratification. In addition, fasting state stratification highlighted aspects of metabolism which manifest themselves uniquely in the postprandial and fasted states. Indeed, while fasted serum has been a source of many markers for metabolic diseases ^36^, individual responses to a meal can carry information regarding metabolic flexibility ^37^, prediabetes state ^38^ or postprandial inflammation ^39^. To our knowledge, the issue of the mixed population of fasted and non-fasted subjects in the biobanked samples has not been previously addressed. As our model was built using absolute quantification it is transferable to other studies, and could be especially useful for cohorts without fasting state information. Of note, the stability of metabolomics factors used to generate the fasting state predictive model during sub-optimal collection practices (i.e. storage at room temperature for days prior to refrigeration) ^40^ and upon prolong freezer storage were previously described ^41^.

The postprandial state is the dominant metabolic state due to the common ingestion of multiple meals yielding 6-8hr postprandial fluctuation in lipoprotein particles ^42^, non-esterified lipids ^43^, hormones ^33^, etc. The strongest positive associations in the non-fasted samples were observed between perceptual speed and levels of non-esterified LA, EPA, DHA, the 15-LOX metabolite of EPA (15-HEPE) and palmitoleate- and EPA-derived ethanolamides (i.e. POEA and EPEA). Other measured ethanolamides did not show significant associations with perceptual speed. The positive association between LA and perceptual speed suggests a role of LA in regulating memory domains, consistent with studies showing reduced LA concentrations in multiple brain regions affected by Alzheimer’s Disease pathology ^44^.

Ethanolamides are generally considered anti-inflammatory ^45^ and neuroprotective ^46^, however, their postprandial physiological consequences are not well understood. Like PUFA, all ethanolamides are lower in non-fasted versus fasted subjects (**Supplemental Fig. S3**) as previously reported^47^. This may suggest that maintaining a higher level of LA and/or POEA and/or EPEA in the postprandial state may reflect metabolism beneficial to perceptual speed and cognition and is not dependent on the “basal” fasted state. The majority of ethanolamide studies have focused on derivatives of AA, oleic acid and palmitic acid, i.e. AEA, OEA and PEA respectively. AEA and PEA can activate CB1 and CB2 receptors ^48^, important players in neuroinflammatory processes ^49^. Moreover, AEA can similarly activate the transient vanilloid receptor type 1 (TRPV1) involved in the transduction of acute and inflammatory pain signals in the periphery ^50^, and have a variety of functions within the central nervous system, and may mediate some excitotoxic effects ^51^. OEA, a peroxisome proliferator-activated receptor α agonist, is a regulator of satiety and sleep with both central and peripheral anorexigenic effects ^48^.

Similarly, a satiety effect was achieved by external administration of the linoleoyl ethanolamide (LEA) and α-linolenoyl ethanolamide (aLEA) respectively ^52^. However, little is known about the biological actions of POEA and EPEA. Additionally, palmitoleic acid and its metabolites are highly abundant in adipose tissue and have been described adipose derived lipokines ^53^, which may indicate a specific involvement of adipose tissue in the maintenance of perceptual speed.

In the non-fasted state, bile acids manifested similar relationships with perceptual speed, semantic and episodic memory and global cognition. Generally, cognitive domains showed positive associations with unconjugated and negative associations with both taurine and glycine conjugated bile acids, the observation strengthened by associations with conjugated/unconjugated bile acid ratios, implying a role for liver metabolism in cognitive maintenance. Of note, the same associations were observed for primary and secondary bile acid. Additionally, we saw negative associations of episodic memory and global cognition with the ratio of both conjugated and unconjugated deoxycholic acid (DCA) to cholic acid (CA) and conjugated lithocholic acid (LCA) to CDCA. Those ratios represent dihydroxylation of primary bile acids (CA and CDCA) by gut bacteria and were previously reported to be negatively associated with cognition ^54^ and atrophy, and brain glucose metabolism in AD ^55^.

These findings suggest increased liver bile acid modification (i.e. conjugation with amino acids), as well as gut microbiome activity may negatively influence cognition. Importantly, these relationships were not observed in fasted samples, suggesting the importance of postprandial metabolism to either drive or highlight these metabolic associations with cognition, warranting further clinical trials using standardized meal tolerance tests.

Using only fasted subjects, we found perceptual speed to be negatively associated with sEH activity reflected by LA-dependent product: substrate ratios ^32^, EPA- and DHA-derived she metabolites, and T-a-MCA and positively associated with the glycine conjugation ratio of UDCA (GUDCA/UDCA). Notably, and the predictive model for perceptual speed depended on both sEH activity assessments and sEH-derived omega 3 diols, these metabolic domains appear to contain independent information. Of note, addition of T-a-MCA provided only slight improvement to the model and in alternate iterations of the model through bootstrapping could be replace by free AA (positively associated with perceptual speed). Therefore, our results implicate eighteen carbon fatty acid metabolism (i.e. sEH action on LA and aLA epoxides) and long chain omega 3 fatty acid metabolism (i.e. sEH activity on EPA and DHA epoxides) in the decline of perceptual speed. This is an agreement with two recent studies which showed negative associations between circulating sEH activity and executive function ^56,57^.

Epoxy fatty acids have potent vasorelaxant and anti-inflammatory properties, while fatty acid diols have demonstrated pro-inflammatory effects and actions as inhibiters of protein kinase B-(i.e. Akt) dependent processes ^58^. Recent studies of mice and men have implicated sEH in neurodegenerative diseases of the brain ^59^. Moreover, DHA feeding enhances the therapeutic efficacy of sEH inhibitors in reducing neurocognitive complications in rodent models of diabetes ^60^. Together, these studies provide strong evidence that the identified shifts in sEH metabolism in association with cognitive decline may be linked to the underlying pathology of this process.

In contrast to the non-fasted state, in the fasted state general association between bile acids metabolism and cognition were not observed, and few specific bile acids showed significant correlations. The ratio of conjugated to unconjugated UDCA was positively associated with episodic memory and global cognition, whereas T-a-MCA was negatively associated with almost all cognitive domains. UDCA and its conjugated derivatives are hydrophilic bile acids previously reported to improve mitochondrial function ^61^ and manifest neuroprotective properties both *in vivo* ^62^ and prevent amyloid-β – induced neuronal death *in vitro* ^63^. T-a-MCA appears in the predictive model for perceptual speed, together with sEH, suggesting their independent association with cognition. GUDCA/UDCA and T-a-MCA both appear in predictive model for episodic memory and global cognition, suggesting their independent associations with cognition.

In conclusion, here we have analyzed serum from the ROS/MAP cohort using a suite of targeted metabolomic assays in search of biomarkers of cognitive function with plausible links to inflammatory responses and energy metabolism. Our study suggests the involvement of sEH and omega-3 PUFA metabolism in cognition. Moreover, during the course of this effort we have produced a tool to determine subject fasting state when unknown, and demonstrated the pivotal nature of this discrimination in biomarker discovery. We have demonstrated that the fasted and non-fasted states carry distinct information regarding the connection of metabolism and cognition. As opportunistically collected non-fasted samples manifest high variance, future studies using a standardized mix meal tolerance test ^64^ could prove useful to validate and discover new relationships between postprandial metabolism and cognition.

## Author Contributions

KB, TP and JWN adapted analytical methods, conducted analyses, and evaluated analytical data quality. KB, AYT and JWN developed statistical analysis plan. KB conducted statistical analyses. DAB obtained study samples. DAB, PLDJ and RK-D were responsible for study design and procured funding. KB and JWN wrote the manuscript. All authors edited and approved the manuscript.

## Acknowledgement

The data presented here are in whole or in part based on data obtained from the AMP-AD Knowledge Portal (https://adknowledgeportal.synapse.org/#/Explore/Data). Metabolomics data is provided by the Alzheimer’s Disease Metabolomics Consortium (ADMC) and funded wholly or in part by the following grants and supplements thereto: NIA R01AG046171, RF1AG051550, RF1AG057452, R01AG059093, RF1AG058942, U01AG061359, U19AG063744 and FNIH: #DAOU16AMPA awarded to Dr. Kaddurah-Daouk at Duke University in partnership with a large number of academic institutions. As such, the investigators within the ADMC, not listed specifically in this publication’s author’s list, provided data but did not participate in analysis or writing of this manuscript. A complete listing of ADMC investigators can be found at: https://sites.duke.edu/adnimetab/team/. The Religious Orders and the Rush Memory and Aging studies (ROS/MAP) was funded by the following NIA grants awarded to Dr. David Bennet at Rush University [P30AG10161, R01AG16819, R01AG17917, U01AG61356]. ROSMAP data can be requested at https://www.radc.rush.edu. Additional support was provided by USDA Intramural Project 2032-51530-022-00D and NIH U24 DK097154 awarded to John W. Newman. The USDA is an equal opportunity employer and provider.

