## Supplemental Figure S1 for "Serum metabolomic biomarkers of perceptual speed in cognitively normal and mildly impaired subjects with fasting state stratification"

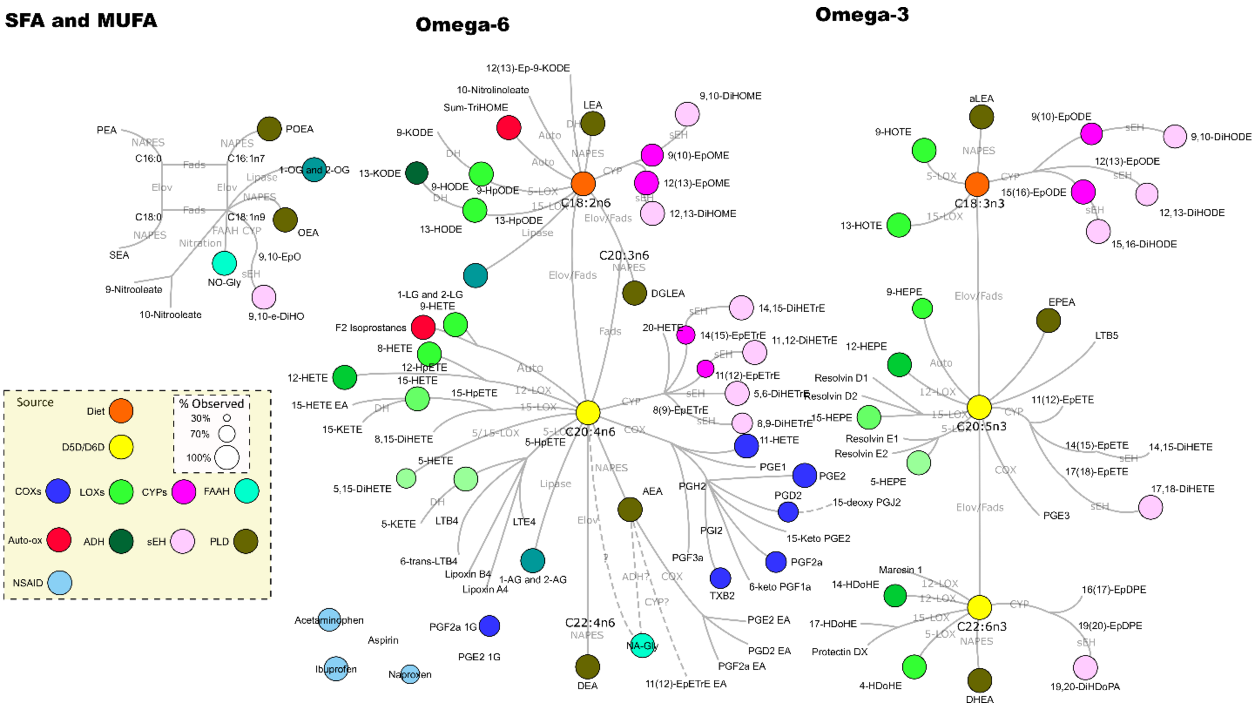


**Supplemental Figure S1.** Oxylipins, endocannabinoids, fatty acids and NSAIDs detected in serum of experimental cohort. Detected metabolites are presented as colored node. Node color represents metabolic pathway/source of metabolite whereas size represents detection frequency. Edge label describe metabolizing enzyme. D5D/D6D – delta 5 and delta 6 desaturase, COXs – cyclooxygenases, LOXs, lipoxygenases, CYPs – cytochrome p450, FAAH – fatty acid amide hydrolase, ADH – alcohol dehydrogenase, sEH – soluble epoxide hydrolase, PLD – phospholipase D, NSAIDs – non-steroidal anti-inflammatory drugs.
