## Supplemental Figure S2 for "Serum metabolomic biomarkers of perceptual speed in cognitively normal and mildly impaired subjects with fasting state stratification"

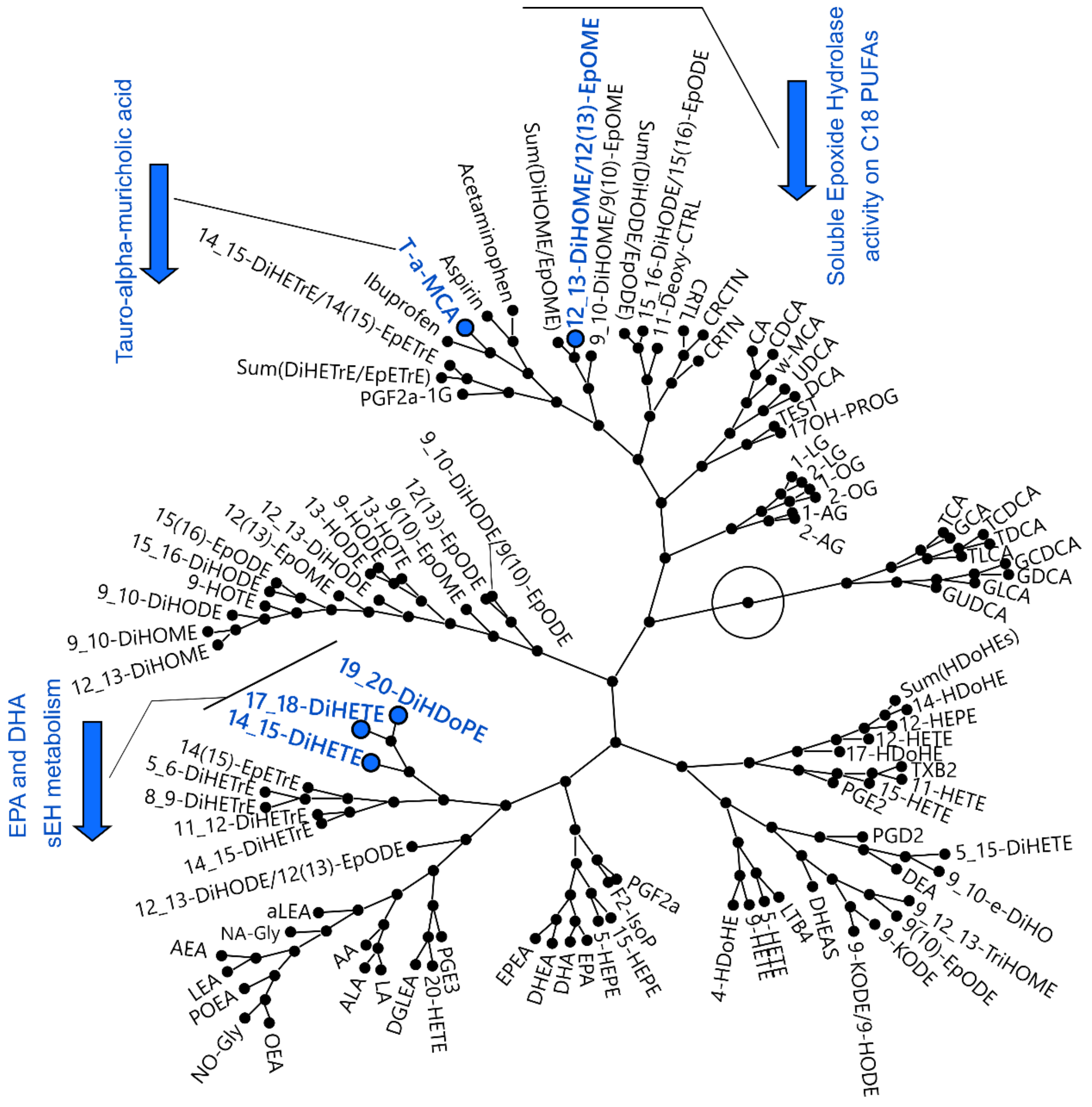


**Supplemental Figure S2**. Constituent metabolites of perceptual speed predictive model, projected onto metabolite correlation network. Constellation plot of hierarchical cluster analysis of oxylipins, endocannabinoids, PUFA and bile acids, recorded in the fasted samples.

Metabolites used un the model are highlighted. Colors indicate directionality of correlation with perceptual speed (blue – negative). General description is provided for metabolites closely correlated with those used in the predictive model.
