## Supplemental Figure S3 for "Serum metabolomic biomarkers of perceptual speed in cognitively normal and mildly impaired subjects with fasting state stratification"

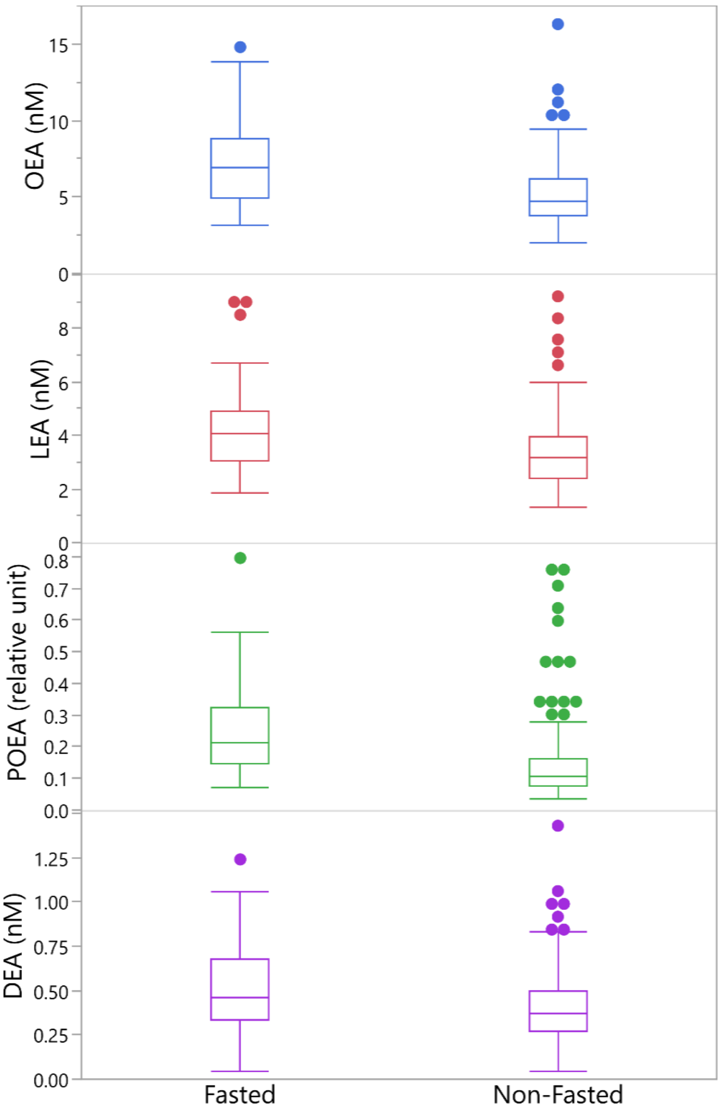


**Supplemental Figure S3**. Difference in the levels of ethanolamides between fasted and non-fasted samples. Mean concentrations of all displayed ethanolamides were different at the $p<0.05$.
