## Supplemental Table S1 for "Serum metabolomic biomarkers of perceptual speed in cognitively normal and mildly impaired subjects with fasting state stratification"

**Supplemental Table S1.** Oxylipins, endocannabinoids, PUFAs, NSAIDs, bile acids and steroids specific information and identifiers.

| Report Order | Analyte-ID | LOD (NM) | LOQ (NM) | Observed | InChIKey | PubChem CID | Chemical class | Enzyme | Parent FA |
|---|---|---|---|---|---|---|---|---|---|
| | | | | | **Oxylipins** | | | | |
| 1 | TXB2 | 0.233 | 0.699 | Yes | XNRNNGPBEPRNAR-JQBLCGNGSA-N | 5283137 | TX | COX1 | AA |
| 2 | 6-keto-PGF1a | 0.253 | 0.759 | No | KFGOFTHODYBSGM-ZUNNJUQCSA-N | 5280888 | PGs | COX1 | AA |
| 3 | PGE1 | 0.568 | 1.7 | No | GMVPRGQOIOIIMI-DWKJAMRDSA-N | 5280723 | PGs | COX2 | DGLA |
| 4 | PGE2 | 0.198 | 0.595 | Yes | XEYBRNLFEZDVAW-ARSRFYASSA-N | 5280360 | PGs | COX2 | AA |
| 5 | 15-Keto-PGE2 | 0.287 | 0.862 | No | YRTJDWROBKPZNV-KMXMBPPJSA-N | 5280719 | PGs | COX2 | AA |
| 6 | PGD2 | 0.267 | 0.802 | Yes | BHMBVRSPMRCCGG-OUTUXVNYSA-N | 448457 | PGs | COX2 | AA |
| 7 | 15-deoxy-PGJ2 | 0.0824 | 0.247 | No | VHRUMKCAEVRUBK-GODQJPCRSA-N | 5311211 | PGs | COX2 | AA |
| 8 | PGF2a | 0.299 | 0.897 | Yes | PXGPLTODNUVGFL-UAAPODJFSA-N | 5283078 | PGs | COX2 | AA |
| 9 | PGE3 | 0.37 | 1.11 | Yes | CBOMORHDRONZRN-QLOYDKTKSA-N | 5280937 | PGs | COX2 | EPA |
| 10 | PGF3a | 0.369 | 1.11 | No | SAKGBZWJAIABSY-SAMSIYEGSA-N | 5280940 | PGs | COX2 | EPA |
| 11 | F2-Isoprostanes | 0.243 | 0.729 | Yes | --- | --- | PGs | Auto-ox | AA |
| 12 | PGE2-EA | 0.264 | 0.792 | No | GKKWUSPPIQURFM-IGDGGSTLSA-N | 5283119 | PG-EA | COX2 | AA |
| 13 | PGF2a-EA | 0.29 | 0.871 | Yes | XCVCLIRZZCGEMU-FPLRWIMGSA-N | 53481911 | PG-EA | COX2 | AA |
| 14 | PGD2-EA | 0.484 | 1.45 | No | KEYDJKSQFDUAGF-YIRKRNQHSA-N | 5283120 | PG-EA | COX2 | AA |
| 15 | PGF2a-1G | 0.468 | 1.4 | Yes | NWKPOVHSHWJQNI-OMVDPNNKSA-N | 24778485 | PG-Gly | COX2 | AA |
| 16 | PGE2-1G | 0.645 | 1.93 | No | RJXVYMMSQBYEHN-SDTVLRMPSA-N | 52193688 | PG-Gly | COX2 | AA |
| 17 | LTE4 | | | No | OTZRAYGBFWZKMX-MPWKMEBCSA-N | 5280749 | Cys-LT | sEH | AA |
| 18 | 6-trans-LTB4 | 0.99 | 2.97 | No | VNYSSYRCGWBHLG-UKNWISKWSA-N | 5283128 | Diol | LOX | AA |
| 19 | LTB4 | 0.217 | 0.652 | Yes | KFGOFTHODYBSGM-ZUNNJUQCSA-N | 5280888 | Diol | LOX | AA |
| 20 | LTB5 | 0.0955 | 0.286 | No | BISQPGCQOHLHQK-HDNPQISLSA-N | 5283125 | Diol | LOX | AA |
| 21 | 8_15-DiHETE | 0.614 | 1.84 | No | NNPWRKSGORGTIM-RCDCWWQHSA-N | 53480358 | Diol | LOX | AA |
| 22 | 5_15-DiHETE | 0.352 | 1.06 | Yes | UXGXCGPWGSUMNI-BVHTXILBSA-N | 5283158 | Diol | LOX | AA |
| 23 | 9_12_13-TriHOME | 0.276 | 0.828 | Yes | MDIUMSLCYIJBQC-MVFSOIOZSA-N | 9858729 | Triol | LOX | LA |
| 24 | Lipoxin-B4 | 0.521 | 1.56 | No | UXVRTOKOJOMENI-WLPVFMORSA-N | 5280915 | Triol | LOX | AA |
| 25 | Resolvin-E1-Screen | | | No | AOPOCGPBAIARAV-WEKRNNBPSA-N | 25063347 | Triol | LOX | AA |
| 26 | Lipoxin-A4 | 0.376 | 1.13 | No | IXAQOQZEOGMIQS-SSQFXEBMSA-N | 5280914 | Triol | LOX | AA |
| 27 | Resolvin-E2-Screen | | | No | KPRHYAOSTOHNQA-NNQKPOSRSA-N | 16061125 | Triol | LOX | AA |
| 28 | Resolvin-D1 | 0.228 | 0.685 | No | OIWTWACQMDFHJG-NJIQAZPPSA-N | 16061135 | Triol | LOX | DHA |
| 29 | Resolvin-D2 | 0.704 | 2.11 | No | IKFAUGXNBOBQDM-XFMPMKITSA-N | 11383310 | Triol | LOX | DHA |
| 30 | Protectin-DX | 0.186 | 0.558 | No | CRDZYJSQHCXHEG-XLBFCUQGSA-N | 11667655 | Diol | LOX | DHA |
| 31 | Maresin-1 | 0.921 | 2.76 | No | HLHYXXBCQOUTGK-LUSCUACYSA-N | 102518309 | Triol | LOX | DHA |
| 32 | 9_10-e-DiHO | 1.15 | 3.45 | Yes | VACHUYIREGFMSP-SJORKVTESA-N | 441460 | vic-Diol | sEH | OA |
| 33 | 12_13-DiHOME | 0.212 | 0.635 | Yes | CQSLTKIXAJTQGA-FLIBITNWSA-N | 10236635 | vic-Diol | sEH | LA |

Continuation of **Table S1**.

| Report Order | Analyte-ID | LOD (NM) | LOQ (NM) | Observed | InChIKey | PubChem CID | Chemical class | Enzyme | Parent FA |
|---|---|---|---|---|---|---|---|---|---|
| 34 | 9_10-DiHOME | 0.206 | 0.617 | Yes | XEBKSQSGNGRGDW-YFHOEESVSA-N | 9966640 | vic-Diol | sEH | LA |
| 35 | 15_16-DiHODE | 0.311 | 0.934 | Yes | LKLLJYJTYPVCID-OHPMOLHNSA-N | 16061068 | vic-Diol | sEH | aLA |
| 36 | 12_13-DiHODE | 0.187 | 0.561 | Yes | RGRKFKRAFZJQMS-OOHFSOINSA-N | 16061067 | vic-Diol | sEH | aLA |
| 37 | 9_10-DiHODE | 0.32 | 0.961 | Yes | QRHSEDZBZMZPOA-ZJSQCTGTSA-N | 16061066 | vic-Diol | sEH | aLA |
| 38 | 14_15-DiHETrE | 0.172 | 0.515 | Yes | SYAWGTIVOGUZMM-ILYOTBPNSA-N | 5283147 | vic-Diol | sEH | AA |
| 39 | 11_12-DiHETrE | 0.112 | 0.336 | Yes | LRPPQRCHCPFBPE-KROJNAHFSA-N | 5283146 | vic-Diol | sEH | AA |
| 40 | 8_9-DiHETrE | 0.298 | 0.895 | Yes | DCJBINATHQHPKO-TYAUOURKSA-N | 5283144 | vic-Diol | sEH | AA |
| 41 | 5_6-DiHETrE | 0.136 | 0.407 | Yes | GFNYAPAJUNPMGH-QNEBEIHSSA-N | 5283142 | vic-Diol | sEH | AA |
| 42 | 14_15-DiHETE | 0.998 | 3 | Yes | BLWCDFIELVFRJY-QXBXTPPVSA-N | 16061119 | vic-Diol | sEH | EPA |
| 43 | 17_18-DiHETE | 1.08 | 3.24 | Yes | XYDVGNAQQFWZEF-JPURVOHMSA-N | 16061120 | vic-Diol | sEH | EPA |
| 44 | 19_20-DiHDoPE | 0.211 | 0.632 | Yes | FFXKPSNQCPNORO-MBYQGORISA-N | 16061148 | vic-Diol | sEH | DHA |
| 45 | 13-HODE | 1.18 | 3.55 | Yes | HNICUWMFWZBIFP-IRQZEAMPSA-N | 6443013 | R-OH | LOX | LA |
| 46 | 9-HODE | 0.645 | 1.94 | Yes | NPDSHTNEKLQQIJ-SIGMCMEVSA-N | 5282945 | R-OH | LOX | LA |
| 47 | 13-HOTE | 0.405 | 1.22 | Yes | KLLGGGQNRTVBSU-JDTPQGGVSA-N | 10469728 | R-OH | LOX | aLA |
| 48 | 9-HOTE | 0.207 | 0.621 | Yes | YUPHIKSLGBATJK-OBKPXJAFSA-N | 53480359 | R-OH | LOX | aLA |
| 49 | 20-HETE | 0.397 | 1.19 | Yes | NNDIXBJHNLFJJP-DTLRTWKJSA-N | 5283157 | R-OH | CYP | AA |
| 50 | 15-HETE | 0.208 | 0.625 | Yes | JSFATNQSLKRBCI-VAEKSGALSA-N | 5280724 | R-OH | LOX | AA |
| 51 | 12-HETE | 0.173 | 0.518 | Yes | ZNHVWPKMFKADKW-FYMOKONMSA-N | 5312983 | R-OH | LOX | AA |
| 52 | 11-HETE | 0.25 | 0.749 | Yes | GCZRCCHPLVMMJE-RSPKXIRXSA-N | 5312981 | R-OH | LOX | AA |
| 53 | 9-HETE | 0.63 | 1.89 | Yes | KATOYYZUTNAWSA-DLJQHUEDSA-N | 5312978 | R-OH | LOX | AA |
| 54 | 8-HETE | 0.651 | 1.95 | Yes | NLUNAYAEIJYXRB-VYOQERLCSA-N | 5283154 | R-OH | LOX | AA |
| 55 | 5-HETE | 0.242 | 0.726 | Yes | KGIJOOYOSFUGPC-JGKLHWIESA-N | 5280733 | R-OH | LOX | AA |
| 56 | 15-HEPE | 0.191 | 0.574 | Yes | UDXLGBLAJBYLSZ-XBCQTNLFSA-N | 53480357 | R-OH | LOX | EPA |
| 57 | 12-HEPE | 0.312 | 0.935 | Yes | MCRJLMXYVFDXLS-QGQBRVLBSA-N | 10041593 | R-OH | LOX | EPA |
| 58 | 9-HEPE | 0.386 | 1.16 | Yes | OXOPDAZWPWFJEW-FPRWAWDYSA-N | 5283187 | R-OH | LOX | EPA |
| 59 | 5-HEPE | 0.706 | 2.12 | Yes | FTAGQROYQYQRHF-FCWZHQICSA-N | 6439678 | R-OH | LOX | EPA |
| 60 | 17-HDoHE | 0.614 | 1.84 | Yes | SWTYBBUBEPPYCX-VIIQGJSXSA-N | 6439179 | R-OH | LOX | DHA |
| 61 | 14-HDoHE | 0.545 | 1.64 | Yes | ZNEBXONKCYFJAF-BGKMTWLOSA-N | 11566378 | R-OH | LOX | DHA |
| 62 | 4-HDoHE | 0.288 | 0.865 | Yes | IFRKCNPQVIJFAQ-JGDWKEERSA-N | 53394255 | R-OH | LOX | DHA |
| 63 | 13-KODE | 1.27 | 3.81 | Yes | JHXAZBBVQSRKJR-BSZOFBHHSA-N | 6446027 | R=0 | ADH | LA |
| 64 | 9-KODE | 1.13 | 3.38 | Yes | LUZSWWYKKLTDHU-ZJHFMPGASA-N | 9839084 | R=0 | ADH | LA |
| 65 | 12(13)-Ep-9-KODE | 0.812 | 2.44 | No | RCMABBHQYMBYKV-BUHFOSPRSA-N | 5283007 | R=0 | ADH | LA |
| 66 | 15-KETE | 0.723 | 2.17 | No | YGJTUEISKATQSM-USWFWKISSA-N | 5280701 | R=0 | ADH | AA |
| 67 | 5-KETE | 3.38 | 10.2 | No | MEASLHGILYBXFO-XTDASVJISA-N | 5283159 | R=0 | ADH | AA |
| 68 | 13-HpODE-Screen | | | No | JDSRHVWSAMTSSN-IRQZEAMPSA-N | 5280720 | R-OOH | LOX | LA |
| 69 | 9-HpODE-Screen | | | No | JGUNZIWGNMQSBM-ZJHFMPGASA-N | 6439847 | R-OOH | LOX | AA |
| 70 | 9(10)-EpO | 2.59 | 7.77 | Yes | IMYZYCNQZDBZBQ-UHFFFAOYSA-N | 15868 | Epox | CYP | OA |
| 71 | 12(13)-EpOME | 0.303 | 0.91 | Yes | CCPPLLJZDQAOHD-FLIBITNWSA-N | 5356421 | Epox | CYP | LA |
| 72 | 9(10)-EpOME | 0.133 | 0.399 | Yes | FBUKMFOXMZRGRB-YFHOEESVSA-N | 6246154 | Epox | CYP | LA |
| 73 | 15(16)-EpODE | 0.498 | 1.49 | Yes | HKSDVVJONLXYKL-OHPMOLHNSA-N | 16061062 | Epox | CYP | aLA |
| 74 | 12(13)-EpODE | 0.407 | 1.22 | Yes | BKKGUKSHPCTUGE-OOHFSOINSA-N | 16061061 | Epox | CYP | aLA |
| 75 | 9(10)-EpODE | 0.443 | 1.33 | Yes | JTEGNNHWOIJBJZ-ZJSQCTGTSA-N | 16061060 | Epox | CYP | aLA |
| 76 | 14(15)-EpETrE | 0.19 | 0.57 | Yes | WLMZMBKVRPUYIG-LTCHCNGXSA-N | 11954058 | Epox | CYP | AA |
| 77 | 11(12)-EpETrE | 0.2 | 0.599 | Yes | DXOYQVHGIODESM-IQCOFVSKSA-N | 53480479 | Epox | CYP | AA |

Continuation of **Table S1**.

| Report Order | Analyte-ID | LOD (NM) | LOQ (NM) | Observed | InChIKey | PubChem CID | Chemical class | Enzyme | Parent FA |
|---|---|---|---|---|---|---|---|---|---|
| 78 | 8(9)-EpETrE | 4.13 | 12.4 | No | DBWQSCSXHFNTMO-TYAUOURKSA-N | 5283203 | Epox | CYP | AA |
| 79 | 17(18)-EpETE | 0.0844 | 0.253 | No | GPQVVJQEBXAKBJ-JPURVOHMSA-N | 16061089 | R-OH | LOX | EPA |
| 80 | 14(15)-EpETE | 0.556 | 1.67 | No | RGZIXZYRGZWDMI-QXBXTPPVSA-N | 16061088 | R-OH | COX | EPA |
| 81 | 11(12)-EpETE | 0.65 | 1.95 | No | QHOKDYBJJBDJGY-BVILWSOJSA-N | 16061087 | Epox | CYP | EPA |
| 82 | 19(20)-EpDoPE | 0.697 | 2.09 | No | OSXOPUBJJDUAOJ-MBYQGORISA-N | 11631565 | Epox | CYP | DHA |
| 83 | 16(17)-EpDoPE | 0.514 | 1.54 | No | BCTXZWCPBLWCRV-ZYADFMMDSA-N | 14392758 | Epox | CYP | DHA |
| 84 | 15-HETE-EA | 0.105 | 0.314 | No | XZQKRCUYLKDPEK-BPVVGZHASA-N | 91886095 | R-OH | LOX | AA |
| 85 | 11(12)-EpETre-EA | 0.06 | 0.18 | No | TYRRSRADDAROSO-KROJNAHFSA-N | 16061183 | Epox | CYP | AA |
| | | | | | **PUFA** | | | | |
| 86 | LA-Screen | | | Yes | OYHQOLUKZRVURQ-HZJYTTRNSA-N | 5280450 | PUFA | Diet | LA |
| 87 | ALA-Screen | | | Yes | DTOSIQBPPRVQHS-PDBXOOCHSA-N | 5280934 | PUFA | Diet | aLA |
| 88 | AA-Screen | | | Yes | YZXBAPSDXZZRGB-DOFZRALJSA-N | 444899 | PUFA | D5D | AA |
| 89 | EPA-Screen | | | Yes | JAZBEHYOTPTENJ-JLNKQSITSA-N | 446284 | PUFA | D6D | EPA |
| 90 | DHA-Screen | | | Yes | MBMBGCFOFBJSGT-KUBAVDMBSA-N | 445580 | PUFA | D6D | DHA |
| | | | | | **Endocannabinoids** | | | | |
| 91 | 10-Nitrolinoleate | 0.381 | 1.14 | No | LELVHAQTWXTCLY-XYWKCAQWSA-N | 5282259 | Nitro-FA | NOS | LA |
| 92 | 9-Nitrooleate | 1.18 | 3.54 | No | CQOAKBVRRVHWKV-UHFFFAOYSA-M | 53412232 | Nitro-FA | NOS | OA |
| 93 | 10-Nitrooleate | 0.565 | 1.7 | No | WRADPCFZZWXOTI-UHFFFAOYSA-N | 53394576 | Nitro-FA | NOS | OA |
| 94 | SEA | 43 | 129 | No | OTGQIQQTPXJQRG-UHFFFAOYSA-N | 27902 | Acyl-EA | PLD | SA |
| 95 | PEA | 4.75 | 14.2 | No | HXYVTAGFYLMHSO-UHFFFAOYSA-N | 4671 | Acyl-EA | PLD | PA |
| 96 | OEA | 0.193 | 0.58 | Yes | BOWVQLFMWHZBEF-KTKRTIGZSA-N | 5283454 | Acyl-EA | PLD | OA |
| 97 | LEA | 0.157 | 0.472 | Yes | KQXDGUVSAAQARU-HZJYTTRNSA-N | 5283446 | Acyl-EA | PLD | LA |
| 98 | aLEA | 0.0796 | 0.239 | Yes | HBJXRRXWHSHZPU-PDBXOOCHSA-N | 5283449 | Acyl-EA | PLD | aLA |
| 99 | DGLEA | 0.149 | 0.447 | Yes | ULQWKETUACYZLI-QNEBEIHSSA-N | 5282272 | Acyl-EA | PLD | DGLA |
| 100 | AEA | 0.133 | 0.399 | Yes | LGEQQWMQCRIYKG-DOFZRALJSA-N | 5281969 | Acyl-EA | PLD | AA |
| 101 | DEA | 0.398 | 1.19 | Yes | FMVHVRYFQIXOAF-DOFZRALJSA-N | 5282273 | Acyl-EA | PLD | Adrenic acid |
| 102 | DHEA | 0.0755 | 0.227 | Yes | CXWASNUDKUTFPQ-KUBAVDMBSA-N | 53245830 | Acyl-EA | PLD | DHA |
| 103 | POEA-Screen | | | Yes | WFRLANWAASSSFV-FPLPWBNLSA-N | 9835868 | Acyl-EA | PLD | POA |
| 104 | EPEA-Screen | | | Yes | OVKKNJPJQKTXIT-JLNKQSITSA-N | 5283450 | Acyl-EA | PLD | EPA |
| 105 | NO-Gly | 0.156 | 0.469 | Yes | HPFXACZRFJDURI-KTKRTIGZSA-N | 6436908 | Acyl-Gly | FAAH | OA |
| 106 | NA-Gly | 0.164 | 0.491 | Yes | YLEARPUNMCCKMP-DOFZRALJSA-N | 5283389 | Acyl-Gly | FAAH | AA |
| 107 | 1-LG | 2.89 | 8.66 | Yes | WECGLUPZRHILCT-GSNKCQISSA-N | 6436630 | MAG | Lipase | LA |
| 108 | 2-LG | 1.2 | 3.61 | Yes | IEPGNWMPIFDNSD-HZJYTTRNSA-N | 5365676 | MAG | Lipase | LG |
| 109 | 1-AG | 0.287 | 0.862 | Yes | DCPCOKIYJYGMDN-HUDVFFLJSA-N | 16019980 | MAG | Lipase | AA |
| 110 | 2-AG | 0.565 | 1.69 | Yes | RCRCTBLIHCHWDZ-DOFZRALJSA-N | 5282280 | MAG | Lipase | AA |
| 111 | 1-OG | 2.67 | 8.01 | Yes | RZRNAYUHWVFMIP-QJRAZLAKSA-N | 12178130 | MAG | Lipase | OA |
| 112 | 2-OG | 1.06 | 3.19 | Yes | UPWGQKDVAURUGE-KTKRTIGZSA-N | 5319879 | MAG | Lipase | OA |
| | | | | | **NSAID** | | | | |
| 113 | Ibuprofen | 2.79 | 8.38 | Yes | HEFNNWSXXWATRW-UHFFFAOYSA-N | 3672 | NSAID | Treatment | NSAID |
| 114 | Aspirin | | | Yes | BSYNRYMUTXBXSQ-UHFFFAOYSA-N | 2244 | NSAID | Treatment | NSAID |
| 115 | Naproxen | 9.07 | 27.2 | Yes | CMWTZPSULFXXJA-VIFPVBQESA-N | 156391 | NSAID | Treatment | NSAID |
| 116 | Acetaminophen | 2.47 | 7.4 | Yes | RZVAJINKPMORJF-UHFFFAOYSA-N | 1983 | NSAID | Treatment | NSAID |

Continuation of **Table S1**.

| Report Order | Analyte ID | LOD (nM | LOQ (nM) | Observed | InChI Key | PubChem_CID | Chemical class | Enzyme | Precursor |
|---|---|---|---|---|---|---|---|---|---|
| **Bile Acids** | | | | | | | | | |
| 1 | CA | 1.41 | 4.23 | No | BHQCQFFYRZLCQQ-OELDTZBJSA-N | 221493 | 1º-BA | Cyp27A1; CYP8B1 | Cholesterol |
| 2 | CDCA | 2.03 | 6.1 | Yes | RUDATBOHQWOJDD-BSWAIDMHSA-N | 10133 | 1º-BA | Cyp27A1 | Cholesterol |
| 3 | UDCA | 1.15 | 3.46 | Yes | RUDATBOHQWOJDD-UZVSRGJWSA-N | 31401 | 2º-BA | Microbiome | CDCA |
| 4 | DCA | 1.36 | 4.08 | Yes | KXGVEGMKQFWNSR-LLQZFEROSA-N | 222528 | 2º-BA | Microbiome | 1º-BA Conj |
| 5 | LCA | 9.43 | 28 | No | SMEROWZSTRWXGI-HVATVPOCSA-N | 9903 | 2º-BA | Microbiome | 1º-BA Conj |
| 6 | w-MCA | 4.97 | 14.9 | Yes | DKPMWHFRUGMUKF-NTPBNISXSA-N | 5283851 | 1º-BA | Cyp27A1 | CDCA |
| 7 | a-MCA | 5.52 | 16.6 | Yes | DKPMWHFRUGMUKF-JDDNAIEOSA-N | 53477700 | 1º-BA | Cyp27A1 | CDCA; b-MCA |
| 8 | b-MCA | 2.64 | 7.91 | Yes | DKPMWHFRUGMUKF-CRKPLTDNSA-N | 5283853 | 1º-BA | Cyp27A1 | a-MCA |
| 9 | TCA | 0.659 | 1.98 | Yes | WBWWGRHZICKQGZ-HZAMXZRMSA-N | 6675 | 1º-BA Conj | BAT | CA |
| 10 | TCDCA | 0.283 | 0.846 | Yes | BHTRKEVKTKCXOH-BJLOMENOSA-N | 387316 | 1º-BA Conj | BAT | CDCA |
| 11 | TDHCA | | | No | UBDJSBRKNHQFPD-PYGYYAGESA-N | 121933 | Pre-1º-BA Conj | BAT | DHCA |
| 12 | TUDCA | 0.047 | 0.143 | Yes | BHTRKEVKTKCXOH-LBSADWJPSA-N | 9848818 | 2º-BA Conj | BAT | UDCA |
| 13 | TDCA | 0.143 | 0.43 | Yes | AWDRATDZQPNJFN-VAYUFCLWSA-N | 2733768 | 2º-BA Conj | BAT | DCA |
| 14 | TLCA | 1.55 | 4.66 | No | QBYUNVOYXHFVKC-GBURMNQMSA-N | 439763 | 2º-BA Conj | BAT | LCA |
| 15 | GCA | 1.38 | 4.17 | Yes | RFDAIACWWDREDC-FRVQLJSFSA-N | 10140 | 1º-BA Conj | BAT | CA |
| 16 | GCDCA | 0.532 | 1.6 | Yes | GHCZAUBVMUEKKP-GYPHWSFCSA-N | 12544 | 1º-BA Conj | BAT | CDCA |
| 17 | GUDCA | 0.621 | 1.86 | Yes | GHCZAUBVMUEKKP-XROMFQGDSA-N | 12310288 | 2º-BA Conj | BAT | UDCA |
| 18 | GDCA | 0.356 | 1.07 | Yes | WVULKSPCQVQLCU-BUXLTGKBSA-N | 3035026 | 2º-BA Conj | BAT | DCA |
| 19 | GHDCA | 0.297 | 0.891 | No | SPOIYSFQOFYOFZ-BRDORRHWSA-N | 114611 | 2º-BA Conj | BAT | HDCA |
| 20 | GLCA | 0.522 | 1.57 | Yes | XBSQTYHEGZTYJE-OETIFKLTSA-N | 115245 | 2º-BA Conj | BAT | LCA |
| 21 | T-w-MCA | 0.505 | 1.52 | No | XSOLDPYUICCHJX-SYCKBGHMSA-N | 118703092 | 1º-BA Conj | BAT | w-MCA |
| 22 | T-a-MCA | 0.505 | 1.52 | Yes | XSOLDPYUICCHJX-QQXJNSDFSA-N | 101657566 | 1º-BA Conj | BAT | a-MCA |
| 23 | T-b-MCA | 0.561 | 1.68 | No | XSOLDPYUICCHJX-UZUDEGBHSA-N | 21124703 | 1º-BA Conj | BAT | b-MCA |

Continuation of the **Table S1.**

| Report Order | Analyte ID | LOD (nM | LOQ (nM) | Observed | InChI Key | PubChem_CID | Chemical class | Enzyme | Precursor |
|---|---|---|---|---|---|---|---|---|---|
| **Steroids** | | | | | | | | | |
| 24 | CRTL | 0.875 | 2.63 | Yes | JYGXADMDTFJGBT-VWUMJDOOSA-N | 5754 | Glucocorticoid | 11-beta-HSD1 & 2; Cyp11B1 | 11-Deoxy-CRTL; CRTN |
| 25 | CRTN | 0.283 | 0.848 | Yes | MFYSYFVPBJMHGN-ZPOLXVRWSA-N | 222786 | Glucocorticoid | 11-beta-HSD 1 & 2 | CRTL |
| 26 | CRCTN | 1.52 | 4.56 | Yes | OMFXVFTZEKFJBZ-HJTSIMOOSA-N | 5753 | Mineralo corticoid | Cyp11B1 | 11-Deoxy-CRTN |
| 27 | 11-Deoxy-CTRL | 0.506 | 1.52 | Yes | WHBHBVVOGNECLV-OBQKJFGGSA-N | 440707 | Glucocorticoid | CYP8B1 | 17OH-Prog; 11-Deoxy-CRTN |
| 28 | E2 | 4.66 | 14 | No | VOXZDWNPVJITMN-ZBRFXRBCSA-N | 5757 | Sex Hormones | Aromatase (CYP19A1) | TEST |
| 30 | TEST | 0.43 | 1.29 | Yes | MUMGGOZAMZWBJJ-DYKIIFRCSA-N | 6013 | Sex Hormones | 3-beta-HSD | 11-Deoxy-CRTL; Andorstenedione |
| 31 | 17OH-Prog | 0.357 | 1.07 | Yes | DBPWSSGDRRHUNT-CEGNMAFCSA-N | 6238 | Corticosteroid | 3-beta-HSD | PROG |
| 32 | PROG | 0.379 | 1.14 | No | RJKFOVLPORLFTN-LEKSSAKUSA-N | 5994 | Sex Hormones | 3-beta-HSD | Pregnenalone |
