## Supplemental Table S2 for "Serum metabolomic biomarkers of perceptual speed in cognitively normal and mildly impaired subjects with fasting state stratification"

**Supplemental Table S2**. Spearman ρ correlation matrix between serum lipid mediators and cognitive domains. Correlation was generated using subjects stratified by the fasting status. Significant correlations (p<0.04) are highlighted in blue. The numbers indicate Spearman ρ. Percent of metabolites manifesting significant correlation with cognitive domains are indicated above the table.

| | | | | | | Fasted | | | | | | Non Fasted | | | | | |
|---|---|---|---|---|---|---|---|---|---|---|---|---|---|---|---|---|---|
| | | | | | **% OxyEndo** | 4% | 2% | 2% | 8% | 2% | 2% | 1% | 1% | 0% | 10% | 2% | 5% |
| | | | | | **% Bile Acids** | 9% | 6% | 6% | 3% | 3% | 0% | 38% | 25% | 0% | 16% | 25% | 0% |
| | | | | | **% steroids** | 0% | 0% | 0% | 0% | 0% | 0% | 0% | 0% | 25% | 0% | 0% | 0% |
| | | | | | **Tatal** | 5% | 3% | 3% | 6% | 2% | 2% | 10% | 7% | 2% | 10% | 8% | 3% |
| **General class** | **Chemical class** | **Enzyme** | **Parent FA** | **Units** | **Variable** | **Episodic memory** | **Global cognition** | **Perceptual orientation** | **Perceptual speed** | **Semantic memory** | **Working memory** | **Spearman ρ Non-Fasted cogn_ep** | **Spearman ρ Non-Fasted cogn_global** | **Spearman ρ Non-Fasted cogn_po** | **Spearman ρ Non-Fasted cogn_ps** | **Spearman ρ Non-Fasted cogn_se** | **Spearman ρ Non-Fasted cogn_wo** |
| OxyEndo | TX | COX1 | AA | nM | TXB2 | 0.017 | -0.054 | 0.0005 | 0.08 | **-0.25** | -0.079 | 0.066 | 0.1 | 0.074 | 0.12 | **0.2** | -0.031 |
| OxyEndo | PGs | COX2 | AA | nM | PGE2 | 0.098 | -0.025 | -0.052 | 0.064 | -0.17 | -0.11 | -0.0052 | 0.03 | -0.06 | 0.084 | 0.15 | 0.026 |
| OxyEndo | PGs | COX2 | AA | nM | PGD2 | 0.14 | 0.16 | 0.12 | **0.25** | 0.088 | 0.021 | 0.0059 | 0.052 | -0.024 | -0.054 | 0.17 | 0.028 |
| OxyEndo | PGs | COX2 | AA | nM | PGF2a | 0.18 | 0.18 | 0.17 | 0.13 | 0.04 | 0.031 | -0.018 | -0.028 | 0.024 | 0.055 | -0.051 | -0.11 |
| OxyEndo | PGs | COX2 | EPA | nM | PGE3 | -0.082 | -0.046 | -0.028 | -0.099 | -0.015 | -0.021 | 0.078 | 0.052 | -0.0075 | -0.053 | -0.0094 | 0.052 |
| OxyEndo | PGs | Auto-ox | AA | nM | F2-IsoP | 0.13 | 0.17 | 0.19 | 0.1 | 0.12 | 0.12 | -0.029 | -0.0034 | 0.039 | 0.087 | -0.074 | -0.047 |
| OxyEndo | PG-Gly | COX2 | AA | nM | PGF2a-1G | 0.03 | -0.024 | **-0.3** | -0.075 | 0.048 | -0.063 | -0.091 | -0.087 | -0.05 | 0.031 | -0.071 | -0.088 |
| OxyEndo | Diol | LOX | AA | nM | LTB4 | **-0.31** | -0.23 | -0.041 | -0.032 | -0.2 | -0.03 | 0.043 | 0.045 | -0.042 | 0.039 | -0.023 | 0.068 |
| OxyEndo | Diol | LOX | AA | nM | 5_15-DiHETE | -0.088 | -0.032 | 0.084 | -0.0091 | -0.028 | 0.06 | -0.0058 | 0.043 | 0.14 | 0.021 | 0.094 | -0.049 |
| OxyEndo | Triol | LOX | LA | nM | 9_12_13-TriHOME | -0.065 | -0.091 | -0.19 | -0.059 | -0.21 | 0.012 | 0.11 | 0.076 | 0.059 | 0.075 | 0.031 | -0.13 |
| OxyEndo | vic-Diol | sEH | OA | nM | 9_10-e-DiHO | -0.063 | 0.017 | -0.019 | 0.061 | -0.04 | 0.13 | 0.077 | 0.06 | -0.029 | 0.11 | 0.074 | -0.045 |
| OxyEndo | vic-Diol | sEH | LA | nM | 12_13-DiHOME | 0.067 | 0.048 | 0.044 | -0.046 | -0.16 | 0.13 | -0.0003 | 0.0075 | 0.04 | 0.033 | -0.025 | -0.015 |
| OxyEndo | vic-Diol | sEH | LA | nM | 9_10-DiHOME | -0.026 | -0.067 | -0.078 | -0.076 | -0.22 | 0.067 | 0.038 | 0.016 | 0.0076 | 0.015 | -0.064 | -0.0086 |
| OxyEndo | vic-Diol | sEH | aLA | nM | 15_16-DiHODE | 0.093 | 0.062 | 0.1 | -0.089 | -0.15 | 0.15 | -0.023 | 0.0061 | 0.0012 | -0.0086 | 0.013 | -0.028 |
| OxyEndo | vic-Diol | sEH | aLA | nM | 12_13-DiHODE | -0.094 | -0.088 | 0.2 | -0.15 | -0.16 | 0.028 | **0.21** | **0.22** | 0.039 | 0.098 | 0.098 | 0.13 |
| OxyEndo | vic-Diol | sEH | aLA | nM | 9_10-DiHODE | 0.0063 | -0.025 | -0.09 | -0.1 | -0.16 | 0.027 | -0.011 | 0.017 | 0.019 | -0.012 | -0.045 | 0.037 |
| OxyEndo | vic-Diol | sEH | AA | nM | 14_15-DiHETrE | 0.032 | 0.023 | -0.14 | -0.09 | -0.04 | 0.16 | -0.013 | 0.0007 | -0.091 | 0.13 | 0.13 | -0.053 |
| OxyEndo | vic-Diol | sEH | AA | nM | 8_9-DiHETrE | -0.085 | -0.097 | -0.091 | -0.13 | -0.21 | 0.069 | -0.031 | -0.033 | -0.059 | 0.041 | 0.055 | -0.069 |
| OxyEndo | vic-Diol | sEH | AA | nM | 5_6-DiHETrE | -0.17 | -0.1 | 0.038 | -0.022 | -0.075 | 0.049 | 0.14 | 0.16 | 0.099 | 0.13 | 0.11 | 0.09 |
| OxyEndo | vic-Diol | sEH | EPA | nM | 14_15-DiHETE | -0.093 | -0.087 | 0.18 | **-0.27** | -0.078 | 0.15 | 0.032 | 0.017 | 0.012 | 0.12 | -0.08 | -0.073 |
| OxyEndo | vic-Diol | sEH | EPA | nM | 17_18-DiHETE | -0.053 | -0.049 | 0.11 | -0.24 | -0.11 | 0.12 | 0.04 | 0.058 | -0.035 | 0.14 | 0.0042 | 0.014 |
| OxyEndo | vic-Diol | sEH | DHA | nM | 19_20-DiHDoPE | -0.17 | -0.18 | 0.12 | **-0.31** | **-0.26** | 0.029 | 0.014 | 0.063 | -0.018 | **0.2** | 0.079 | 0.0052 |
| OxyEndo | R-OH | LOX | LA | nM | 13-HODE | 0.099 | 0.037 | -0.0097 | -0.064 | -0.23 | 0.11 | -0.0029 | 0.01 | 0.038 | 0.11 | -0.02 | -0.019 |
| OxyEndo | R-OH | LOX | LA | nM | 9-HODE | 0.12 | 0.035 | -0.078 | -0.035 | -0.18 | 0.073 | 0.016 | 0.036 | -0.0021 | 0.12 | 0.017 | 0.015 |
| OxyEndo | R-OH | LOX | aLA | nM | 13-HOTE | 0.08 | 0.073 | 0.055 | -0.017 | -0.1 | 0.16 | -0.028 | 0.025 | 0.066 | 0.088 | 0.021 | -0.0026 |
| OxyEndo | R-OH | LOX | aLA | nM | 9-HOTE | 0.12 | 0.081 | 0.041 | -0.085 | -0.12 | 0.13 | 0.05 | 0.078 | 0.0089 | 0.1 | 0.066 | 0.01 |
| OxyEndo | R-OH | CYP | AA | nM | 20-HETE | -0.1 | 0.053 | 0.0078 | 0.11 | 0.18 | -0.011 | -0.029 | -0.073 | 0.0091 | 0.0075 | 0.026 | **-0.2** |
| OxyEndo | R-OH | LOX | AA | nM | 15-HETE | 0.019 | -0.031 | -0.026 | -0.022 | -0.16 | -0.026 | -0.027 | -0.036 | 0.015 | 0.017 | 0.0014 | -0.023 |
| OxyEndo | R-OH | LOX | AA | nM | 12-HETE | 0.0067 | -0.049 | 0.039 | -0.09 | -0.21 | 0.0017 | 0.0035 | -0.023 | -0.019 | -0.043 | 0.028 | -0.0046 |
| OxyEndo | R-OH | LOX | AA | nM | 11-HETE | 0.15 | 0.059 | -0.073 | 0.059 | -0.14 | 0.011 | -0.022 | -0.022 | 0.042 | 0.0063 | 0.077 | -0.062 |
| OxyEndo | R-OH | LOX | AA | nM | 9-HETE | -0.044 | -0.08 | -0.21 | -0.0013 | -0.14 | -0.01 | -0.089 | -0.066 | -0.053 | 0.11 | -0.028 | -0.044 |
| OxyEndo | R-OH | LOX | AA | nM | 5-HETE | -0.13 | -0.091 | -0.13 | 0.11 | -0.025 | 0.0046 | -0.036 | -0.038 | 0.053 | 0.046 | -0.064 | -0.021 |
| OxyEndo | R-OH | LOX | EPA | nM | 15-HEPE | 0.097 | 0.16 | 0.14 | 0.023 | 0.073 | 0.12 | 0.09 | 0.12 | 0.1 | **0.2** | 0.051 | 0.011 |
| OxyEndo | R-OH | LOX | EPA | nM | 12-HEPE | 0.062 | 0.0017 | 0.11 | -0.15 | -0.21 | 0.038 | -0.019 | -0.03 | -0.011 | 0.024 | -0.011 | -0.019 |
| OxyEndo | R-OH | LOX | EPA | nM | 5-HEPE | 0.068 | 0.035 | 0.11 | -0.084 | -0.0083 | -0.017 | -0.0053 | 0.0004 | 0.11 | 0.16 | -0.053 | -0.13 |
| OxyEndo | R-OH | LOX | DHA | nM | 17-HDoHE | -0.053 | -0.091 | 0.097 | -0.18 | -0.21 | -0.0028 | -0.038 | -0.032 | 0.027 | -0.018 | 0.009 | -0.019 |
| OxyEndo | R-OH | LOX | DHA | nM | 14-HDoHE | -0.0074 | -0.038 | 0.097 | -0.12 | -0.21 | 0.033 | -0.018 | -0.037 | 0.025 | 0.015 | -0.014 | -0.041 |
| OxyEndo | R-OH | LOX | DHA | nM | 4-HDoHE | -0.014 | -0.05 | 0.034 | -0.15 | -0.069 | 0.0064 | 0.073 | 0.12 | 0.0038 | **0.18** | 0.032 | 0.079 |
| OxyEndo | R=0 | ADH | LA | nM | 9-KODE | -0.019 | 0.013 | -0.0085 | 0.067 | -0.041 | 0.11 | 0.02 | 0.025 | -0.11 | 0.13 | 0.05 | -0.021 |
| OxyEndo | Epox | CYP | LA | nM | 12(13)-EpOME | 0.078 | 0.14 | 0.099 | 0.19 | -0.012 | 0.17 | 0.059 | 0.11 | 0.058 | 0.11 | 0.031 | 0.1 |
| OxyEndo | Epox | CYP | LA | nM | 9(10)-EpOME | 0.018 | 0.012 | 0.06 | 0.017 | -0.13 | 0.13 | 0.05 | 0.068 | 0.074 | 0.13 | -0.016 | 0.044 |
| OxyEndo | Epox | CYP | aLA | nM | 15(16)-EpODE | **0.27** | **0.28** | 0.15 | 0.057 | 0.017 | **0.33** | -0.013 | 0.023 | -0.054 | 0.074 | 0.021 | 0.041 |
| OxyEndo | Epox | CYP | aLA | nM | 12(13)-EpODE | -0.13 | -0.021 | 0.04 | -0.075 | -0.053 | 0.15 | 0.073 | 0.13 | -0.033 | 0.064 | **0.18** | 0.065 |
| OxyEndo | Epox | CYP | aLA | nM | 9(10)-EpODE | -0.06 | -0.048 | 0.015 | 0.034 | -0.095 | -0.035 | -0.0065 | -0.045 | -0.07 | 0.034 | -0.083 | -0.052 |
| OxyEndo | Epox | CYP | AA | nM | 14(15)-EpETrE | 0.053 | 0.08 | 0.015 | 0.13 | 0.062 | 0.12 | 0.0082 | 0.046 | 0.069 | 0.097 | 0.12 | -0.012 |
| OxyEndo | PUFA | Diet | LA | Rel Abs | LA_screen | 0.005 | 0.11 | 0.016 | 0.1 | 0.048 | 0.2 | -0.03 | 0.0014 | -0.042 | **0.25** | 0.1 | -0.1 |
| OxyEndo | PUFA | Diet | aLA | Rel Abs | ALA_screen | 0.087 | 0.14 | 0.07 | 0.031 | 0.024 | 0.12 | -0.029 | -0.015 | -0.02 | 0.17 | 0.052 | -0.1 |
| OxyEndo | PUFA | D5D | AA | Rel Abs | AA_screen | 0.11 | 0.19 | 0.13 | **0.26** | 0.13 | 0.12 | -0.037 | -0.09 | -0.019 | 0.1 | 0.013 | **-0.21** |
| OxyEndo | PUFA | D6D | EPA | Rel Abs | EPA_screen | 0.2 | 0.2 | 0.22 | 0.025 | 0.064 | 0.099 | -0.053 | -0.0059 | 0.037 | **0.22** | 0.058 | -0.14 |
| OxyEndo | PUFA | D6D | DHA | Rel Abs | DHA_screen | 0.077 | 0.12 | **0.25** | 0.028 | 0.036 | 0.036 | -0.016 | 0.026 | 0.075 | **0.25** | 0.077 | -0.14 |
| OxyEndo | Acyl-EA | PLD | OA | nM | OEA | -0.069 | -0.099 | -0.035 | -0.16 | -0.21 | 0.012 | 0.034 | 0.03 | 0.05 | 0.1 | 0.091 | -0.11 |
| OxyEndo | Acyl-EA | PLD | LA | nM | LEA | -0.064 | -0.11 | -0.13 | -0.13 | -0.1 | 0.0016 | -0.019 | -0.023 | -0.06 | 0.046 | 0.034 | -0.03 |
| OxyEndo | Acyl-EA | PLD | aLA | nM | aLEA | -0.1 | -0.14 | 0.039 | -0.21 | -0.12 | -0.063 | 0.041 | 0.067 | -0.02 | 0.061 | 0.12 | 0.011 |
| OxyEndo | Acyl-EA | PLD | DGLA | nM | DGLEA | -0.15 | -0.1 | -0.23 | -0.073 | -0.06 | 0.033 | -0.02 | 0.031 | 0.0076 | 0.16 | 0.089 | 0.031 |
| OxyEndo | Acyl-EA | PLD | AA | nM | AEA | -0.098 | -0.062 | -0.045 | -0.064 | -0.096 | 0.086 | -0.11 | -0.097 | -0.015 | 0.071 | -0.064 | -0.049 |
| OxyEndo | Acyl-EA | PLD | Adrenic acid | nM | DEA | -0.15 | -0.076 | -0.095 | 0.077 | -0.073 | 0.062 | 0.036 | -0.077 | -0.035 | -0.0032 | -0.039 | **-0.21** |
| OxyEndo | Acyl-EA | PLD | DHA | nM | DHEA | -0.027 | -0.04 | 0.11 | -0.18 | -0.026 | -0.027 | 0.061 | 0.045 | 0.018 | 0.17 | -0.032 | -0.043 |
| OxyEndo | Acyl-EA | PLD | POA | Rel Abs | POEA_Screen | -0.11 | -0.058 | -0.016 | -0.13 | -0.075 | 0.12 | 0.018 | 0.045 | 0.067 | **0.24** | 0.032 | -0.075 |
| OxyEndo | Acyl-EA | PLD | EPA | Rel Abs | EPEA_Screen | 0.053 | 0.071 | -0.0096 | -0.12 | 0.083 | 0.039 | -0.038 | -0.018 | 0.1 | **0.18** | -0.036 | -0.13 |
| OxyEndo | Acyl-Gly | FAAH | OA | nM | NO-Gly | -0.079 | -0.1 | -0.0008 | -0.12 | -0.15 | -0.084 | 0.013 | 0.012 | -0.034 | 0.13 | 0.1 | -0.052 |
| OxyEndo | Acyl-Gly | FAAH | AA | nM | NA-Gly | 0.14 | 0.16 | -0.028 | 0.23 | 0.14 | 0.098 | -0.074 | -0.087 | 0.098 | 0.051 | -0.022 | -0.13 |
| OxyEndo | MAG | Lipase | LA | nM | 1-LG | 0.078 | 0.08 | -0.074 | 0.13 | 0.053 | -0.021 | -0.0075 | -0.067 | -0.055 | 0.016 | -0.03 | -0.15 |
| OxyEndo | MAG | Lipase | LG | nM | 2-LG | 0.069 | 0.057 | -0.077 | 0.1 | 0.05 | -0.034 | -0.034 | -0.096 | -0.056 | -0.012 | -0.052 | -0.16 |
| OxyEndo | MAG | Lipase | AA | nM | 1-AG | 0.1 | 0.082 | -0.082 | 0.16 | 0.062 | -0.067 | -0.067 | -0.13 | -0.043 | -0.054 | -0.066 | -0.15 |
| OxyEndo | MAG | Lipase | AA | nM | 2-AG | 0.11 | 0.091 | -0.082 | 0.19 | 0.078 | -0.099 | -0.096 | -0.15 | -0.028 | -0.05 | -0.074 | **-0.18** |
| OxyEndo | MAG | Lipase | OA | nM | 1-OG | 0.063 | 0.055 | -0.069 | 0.12 | 0.043 | -0.049 | -0.014 | -0.045 | -0.01 | 0.065 | 0.0006 | -0.15 |
| OxyEndo | MAG | Lipase | OA | nM | 2-OG | 0.09 | 0.079 | 0.0049 | 0.1 | 0.014 | -0.023 | 0.0024 | -0.035 | -0.018 | 0.0097 | 0.012 | -0.1 |
| OxyEndo | - | - | - | nM | Sum_n3_Diols | -0.11 | -0.11 | 0.11 | **-0.28** | -0.17 | 0.1 | 0.023 | 0.048 | -0.036 | 0.16 | 0.018 | -0.0041 |
| OxyEndo | - | - | - | nM | Sum(DiHETEs) | -0.052 | -0.054 | 0.12 | **-0.25** | -0.12 | 0.14 | 0.029 | 0.049 | -0.031 | 0.15 | 0.0064 | -0.0061 |
| OxyEndo | - | - | - | nM | Sum(HDoHEs) | -0.018 | -0.051 | 0.087 | -0.13 | -0.22 | 0.028 | -0.012 | -0.034 | 0.024 | 0.023 | -0.012 | -0.047 |
| OxyEndo | - | - | - | Ratio | 15_16-DiHODE/15(16)-EpODE | **-0.27** | **-0.32** | -0.066 | -0.17 | -0.2 | **-0.28** | 0.034 | 0.05 | 0.11 | -0.045 | 0.07 | -0.044 |
| OxyEndo | - | - | - | Ratio | 12_13-DiHOME/12(13)-EpOME | -0.051 | -0.16 | -0.076 | **-0.32** | -0.14 | -0.16 | -0.11 | -0.11 | -0.025 | -0.018 | 0.019 | -0.15 |
| OxyEndo | - | - | - | Ratio | 14_15-DiHETrE/14(15)-EpETrE | -0.055 | -0.099 | -0.075 | -0.24 | -0.12 | -0.071 | -0.0092 | -0.052 | -0.084 | -0.027 | -0.084 | -0.043 |
| OxyEndo | - | - | - | Ratio | 11_12-DiHETrE | -0.024 | 0.0095 | -0.093 | -0.075 | -0.086 | 0.22 | -0.033 | 0.012 | 0.023 | 0.13 | 0.14 | -0.053 |
| OxyEndo | - | - | - | Ratio | 12-HEPE/12-HETE | 0.13 | 0.13 | 0.17 | -0.14 | 0.028 | 0.072 | -0.064 | -0.0078 | 0.009 | 0.15 | -0.077 | 0.039 |
| OxyEndo | - | - | - | Ratio | 9,10-DiHOME/EpOME 2 | 0.054 | 0.059 | -0.13 | 0.018 | 0.11 | 0.006 | -0.029 | -0.095 | -0.088 | -0.12 | -0.044 | -0.15 |
| OxyEndo | - | - | - | Ratio | 9_10-DiHOME/9(10)-EpOME | 0.054 | 0.059 | -0.13 | 0.018 | 0.11 | 0.006 | -0.029 | -0.095 | -0.088 | -0.12 | -0.044 | -0.15 |
| OxyEndo | - | - | - | Ratio | Sum(DiHODE/EpODE) | -0.16 | -0.24 | -0.019 | -0.14 | -0.17 | -0.23 | 0.0027 | 0.036 | 0.11 | -0.043 | 0.064 | -0.032 |
| OxyEndo | - | - | - | Ratio | 9-KODE/9-HODE | -0.069 | 0.039 | 0.09 | 0.078 | 0.15 | 0.11 | 0.012 | 0.015 | -0.095 | 0.033 | 0.061 | -0.021 |
| OxyEndo | - | - | - | Ratio | Sum(DiHETrE/EpETrE) | -0.093 | -0.13 | -0.076 | -0.22 | -0.14 | -0.092 | 0.02 | -0.015 | -0.056 | -0.031 | -0.056 | -0.0044 |
| OxyEndo | - | - | - | Ratio | 12_13-DiHODE/12(13)-EpODE | 0.057 | -0.0019 | 0.14 | 0.045 | -0.074 | -0.054 | 0.17 | 0.13 | 0.11 | 0.03 | -0.067 | 0.12 |
| OxyEndo | - | - | - | Ratio | 14,15/11,12-DiHETrE | 0.051 | -0.056 | -0.1 | -0.11 | 0.031 | -0.24 | 0.078 | 0.036 | -0.12 | 0.037 | -0.002 | 0.0049 |

Continuation of **Supplemental Table S2.**

| General class | Chemical class | Enzyme | Parent FA | Units | Variable | Episodic memory | Global cognition | Perceptual orientation | Perceptual speed | Semantic memory | Working memory | Spearman ρ Non-Fasted cogn_ep | Spearman ρ Non-Fasted cogn_global | Spearman ρ Non-Fasted cogn_po | Spearman ρ Non-Fasted cogn_ps | Spearman ρ Non-Fasted cogn_se | Spearman ρ Non-Fasted cogn_wo |
|---|---|---|---|---|---|---|---|---|---|---|---|---|---|---|---|---|---|
| OxyEndo | - | - | - | Ratio | 9_10-DiHODE/9(10)-EpODE | 0.1 | 0.024 | -0.056 | -0.084 | -0.041 | -0.04 | -0.066 | 0.015 | 0.069 | 0.0007 | 0.04 | 0.047 |
| NSAD | NSAID | Treatment | | nM | Ibuprofen | -0.13 | -0.14 | -0.15 | 0.0081 | -0.056 | -0.19 | -0.1 | -0.11 | -0.012 | -0.0086 | -0.05 | -0.13 |
| NSAD | NSAID | Treatment | | nM | Aspirin | 0.044 | -0.0008 | 0.081 | 0.0069 | -0.026 | -0.12 | 0.018 | 0.051 | 0.023 | 0.085 | -0.055 | 0.13 |
| NSAD | NSAID | Treatment | | nM | Acetaminophen | -0.052 | -0.13 | -0.024 | -0.046 | -0.16 | -0.1 | 0.0038 | -0.083 | -0.021 | -0.14 | -0.14 | -0.042 |
| Bile Acid | Cyp27A1; CYP8B1 | 1º-BA | - | nM | CA | 0.21 | 0.17 | 0.16 | 0.089 | 0.057 | -0.022 | 0.15 | 0.096 | -0.11 | 0.14 | 0.14 | -0.093 |
| Bile Acid | Cholesterol | Cyp27A1 | - | nM | CDCA | **0.27** | 0.2 | 0.061 | 0.11 | 0.11 | -0.047 | 0.094 | 0.15 | -0.0068 | **0.2** | **0.19** | 0.043 |
| Bile Acid | CDCA | Microbiome | - | nM | UDCA | -0.023 | -0.096 | **-0.25** | -0.067 | -0.013 | -0.13 | -0.083 | -0.11 | 0.02 | -0.12 | -0.047 | -0.078 |
| Bile Acid | 1º-BA Conj | Microbiome | - | nM | DCA | 0.2 | 0.17 | -0.086 | 0.19 | 0.21 | -0.029 | 0.014 | 0.12 | 0.11 | 0.078 | **0.2** | 0.1 |
| Bile Acid | CDCA | Cyp27A1 | - | nM | w-MCA | 0.091 | 0.068 | -0.029 | 0.07 | 0.15 | -0.028 | 0.1 | 0.14 | 0.081 | 0.066 | 0.15 | 0.063 |
| Bile Acid | CA | BAT | - | nM | TCA | 0.071 | 0.048 | 0.08 | -0.04 | -0.05 | 0.15 | -0.086 | -0.096 | -0.04 | -0.026 | -0.11 | -0.046 |
| Bile Acid | CDCA | BAT | - | nM | TCDCA | **0.058** | 0.006 | 0.052 | -0.1 | -0.071 | 0.11 | -0.075 | -0.14 | -0.1 | -0.084 | **-0.2** | -0.1 |
| Bile Acid | DCA | BAT | - | nM | TDCA | -0.035 | -0.044 | 0.014 | -0.013 | -0.03 | 0.031 | **-0.18** | -0.14 | 0.0071 | -0.09 | -0.12 | -0.016 |
| Bile Acid | BAT | 2º-BA Conj | - | nM | TLCA | 0.013 | -0.033 | -0.023 | -0.072 | -0.11 | 0.067 | **-0.28** | **-0.29** | -0.086 | **-0.2** | **-0.27** | -0.059 |
| Bile Acid | CA | BAT | - | nM | GCA | 0.18 | 0.15 | 0.13 | 0.018 | 0.1 | 0.11 | -0.11 | -0.073 | -0.052 | 0.027 | -0.051 | -0.019 |
| Bile Acid | CDCA | BAT | - | nM | GCDCA | 0.22 | 0.19 | 0.057 | 0.021 | 0.18 | 0.098 | -0.05 | -0.082 | -0.073 | -0.025 | -0.13 | -0.04 |
| Bile Acid | UDCA | BAT | - | nM | GUDCA | 0.21 | 0.17 | 0.0077 | 0.015 | 0.22 | 0.053 | -0.02 | -0.045 | -0.15 | -0.056 | -0.014 | 0.0001 |
| Bile Acid | DCA | BAT | - | nM | GDCA | 0.15 | 0.14 | -0.011 | 0.15 | 0.16 | 0.045 | **-0.21** | -0.13 | 0.0098 | -0.08 | -0.051 | 0.021 |
| Bile Acid | LCA | BAT | - | nM | GLCA | 0.11 | 0.063 | -0.017 | -0.012 | 0.059 | -0.024 | -0.13 | -0.056 | -0.037 | -0.062 | 0.016 | 0.071 |
| Bile Acid | BAT | 1º-BA Conj | - | nM | T-a-MCA | **-0.29** | **-0.31** | -0.043 | **-0.28** | **-0.26** | -0.066 | -0.081 | -0.14 | -0.067 | -0.16 | -0.15 | 0.0093 |
| Bile Acid | - | - | - | Ratio | GUDCA/UDCA | **0.3** | **0.32** | **0.33** | 0.081 | 0.24 | 0.2 | 0.047 | 0.049 | -0.12 | 0.072 | 0.0068 | 0.052 |
| Bile Acid | - | - | - | Ratio | GDCA/CA | -0.11 | -0.077 | -0.14 | -0.0008 | 0.045 | 0.05 | **-0.28** | **-0.18** | 0.089 | -0.16 | -0.13 | 0.072 |
| Bile Acid | - | - | - | Ratio | TDCA/CA | -0.17 | -0.15 | -0.13 | -0.084 | -0.049 | 0.05 | **-0.26** | **-0.19** | 0.048 | -0.15 | -0.17 | 0.027 |
| Bile Acid | - | - | - | Ratio | TLCA/CDCA | -0.16 | -0.14 | -0.085 | -0.13 | -0.14 | 0.088 | **-0.24** | **-0.27** | -0.025 | **-0.24** | **-0.28** | -0.057 |
| Bile Acid | - | - | - | Ratio | GDCA/DCA | -0.038 | -0.047 | 0.12 | -0.097 | -0.093 | 0.042 | **-0.23** | **-0.27** | -0.14 | -0.16 | **-0.3** | -0.062 |
| Bile Acid | - | - | - | Ratio | GCA/CA | -0.062 | -0.057 | -0.053 | -0.091 | 0.013 | 0.093 | **-0.22** | -0.15 | 0.04 | -0.092 | -0.13 | 0.075 |
| Bile Acid | - | - | - | Ratio | TCA/CA | -0.1 | -0.11 | -0.041 | -0.11 | -0.084 | 0.071 | **-0.21** | -0.16 | 0.021 | -0.099 | -0.16 | 0.018 |
| Bile Acid | - | - | - | Ratio | DCA/CA | -0.034 | 0.011 | -0.2 | 0.074 | 0.16 | 0.091 | **-0.19** | -0.062 | 0.15 | -0.089 | -0.0091 | 0.13 |
| Bile Acid | - | - | - | Ratio | TDCA/DCA | -0.14 | -0.17 | 0.049 | -0.19 | -0.23 | 0.053 | **-0.18** | **-0.22** | -0.08 | -0.12 | **-0.25** | -0.081 |
| Bile Acid | - | - | - | Ratio | (GDCA+GLCA)/(TDCA+TLCA) | 0.2 | 0.16 | 0.022 | -0.063 | 0.15 | 0.017 | **0.18** | **0.19** | 0.011 | 0.14 | 0.15 | 0.098 |
| Bile Acid | - | - | - | Ratio | GCDCA/CDCA | -0.14 | -0.11 | -0.0041 | -0.22 | -0.093 | 0.18 | -0.13 | **-0.2** | -0.034 | **-0.24** | **-0.28** | -0.054 |
| Bile Acid | - | - | - | Ratio | GLCA/CDCA | -0.17 | -0.16 | -0.11 | -0.16 | -0.065 | 0.032 | -0.16 | -0.14 | 0.022 | **-0.19** | -0.14 | 0.027 |
| Bile Acid | - | - | - | Ratio | TDCA/TLCA | -0.029 | -0.053 | 0.0025 | 0.055 | 0.012 | -0.1 | 0.064 | 0.14 | 0.075 | 0.1 | 0.13 | 0.095 |
| Bile Acid | - | - | - | Ratio | GCA/GDCA | 0.077 | 0.058 | 0.2 | -0.09 | -0.06 | 0.1 | 0.06 | 0.006 | -0.11 | 0.094 | -0.028 | -0.061 |
| Bile Acid | - | - | - | Ratio | CA/CDCA | -0.065 | -0.053 | 0.16 | -0.034 | -0.15 | -0.023 | 0.11 | -0.016 | -0.12 | -0.0093 | 0.0069 | -0.15 |
| Bile Acid | - | - | - | Ratio | GCDCA/GLCA | 0.03 | 0.033 | 0.083 | -0.0095 | -0.023 | 0.091 | 0.066 | -0.012 | -0.0069 | 0.028 | -0.09 | -0.09 |
| Bile Acid | - | - | - | Ratio | GDCA/GLCA | 0.061 | 0.064 | -0.021 | 0.13 | 0.02 | 0.11 | -0.068 | -0.029 | 0.038 | 0.012 | -0.02 | 0.033 |
| Steroids | 11-Deoxy-CRTL; CRTN | 11-beta-HSD; Cyp11B1 | - | nM | CRTL | -0.021 | -0.0032 | -0.046 | -0.052 | 0.15 | 0.021 | -0.0089 | -0.07 | -0.071 | -0.066 | -0.023 | -0.067 |
| Steroids | CRTL | 11-beta-HSD | - | nM | CRTN | -0.023 | -0.012 | -0.016 | -0.12 | 0.024 | 0.11 | -0.015 | -0.062 | -0.082 | -0.071 | -0.038 | -0.025 |
| Steroids | 11-Deoxy-CRTN | Cyp11B1 | - | nM | CRCTN | -0.044 | -0.06 | -0.054 | -0.13 | 0.14 | -0.078 | -0.0018 | -0.051 | -0.055 | -0.076 | 0.012 | -0.048 |
| Steroids | 17OH-Prog; 11-Deoxy-CRTN | CYP8B1 | - | nM | 11-Deoxy-CTRL | -0.073 | -0.016 | 0.099 | 0.087 | 0.059 | 0.088 | 0.019 | -0.052 | -0.065 | -0.13 | -0.051 | -0.11 |
| Steroids | - | - | - | nM | DHEAS | -0.061 | -0.01 | 0.0026 | 0.0047 | 0.17 | 0.067 | 0.02 | 0.032 | 0.0046 | 0.093 | 0.019 | -0.055 |
| Steroids | 11-Deoxy-CRTL; Andorstenedione | 3-beta-HSD | - | nM | TEST | 0.024 | 0.074 | -0.011 | 0.039 | -0.023 | 0.23 | 0.053 | 0.025 | -0.012 | -0.014 | -0.0075 | -0.024 |
| Steroids | PROG | 3-beta-HSD | - | nM | 17OH-PROG | 0.018 | 0.035 | 0.092 | 0.013 | 0.05 | 0.11 | 0.054 | 0.017 | **-0.2** | 0.016 | 0.027 | -0.0049 |
| Steroids | - | - | - | Ratio | Tes/Prog | -0.041 | 0.0086 | -0.11 | 0.028 | -0.084 | 0.18 | 0.036 | 0.021 | **0.18** | -0.028 | -0.042 | -0.053 |
