## Supplemental Table S3 for "Serum metabolomic biomarkers of perceptual speed in cognitively normal and mildly impaired subjects with fasting state stratification"

**Supplemental Table S4**. Spearman's ρ correlation between cognitive domains and lipid mediators in all subjects (without fasting state stratification).

| | Spearman ρ | | | | | | Correlation p-value | | | | | |
|---|---|---|---|---|---|---|---|---|---|---|---|---|
| Meatbolite | Episodic memory | Global cognition | Perceptual orientation | perceptual speed | semantic memory | working memmory | Episodic memory | Global cognition | Perceptual orientation | perceptual speed | semantic memory | working memmory |
| T-a-MCA/CDCA | -0.218 | -0.246 | -0.0716 | -0.23 | -0.245 | -0.0175 | 0.0015 | 0.0003 | 0.302 | 0.0008 | 0.0003 | 0.801 |
| GCDCA/CDCA | -0.141 | -0.175 | -0.059 | -0.204 | -0.196 | 0.0205 | 0.041 | 0.0113 | 0.395 | 0.003 | 0.0044 | 0.768 |
| LA_screen | 0.0101 | 0.0511 | 0.0104 | 0.201 | 0.0613 | -0.0111 | 0.884 | 0.46 | 0.88 | 0.0034 | 0.376 | 0.873 |
| T-a-MCA | -0.158 | -0.196 | -0.0653 | -0.199 | -0.187 | -0.0145 | 0.0219 | 0.0044 | 0.347 | 0.0038 | 0.0066 | 0.835 |
| TLCA/CDCA | -0.212 | -0.228 | -0.0637 | -0.19 | -0.225 | -0.0072 | 0.002 | 0.0009 | 0.359 | 0.0058 | 0.001 | 0.918 |
| DHA_screen | 0.0452 | 0.075 | 0.142 | 0.187 | 0.0557 | -0.0666 | 0.514 | 0.278 | 0.0393 | 0.0066 | 0.421 | 0.336 |
| w-MCA/T-a-MCA | 0.196 | 0.223 | 0.0839 | 0.186 | 0.234 | 0.003 | 0.0043 | 0.0012 | 0.226 | 0.007 | 0.0006 | 0.966 |
| GLCA/CDCA | -0.179 | -0.158 | -0.0462 | -0.17 | -0.101 | 0.0352 | 0.0092 | 0.0219 | 0.505 | 0.0136 | 0.147 | 0.612 |
| AA_screen | 0.0432 | 0.0237 | 0.0507 | 0.167 | 0.032 | -0.099 | 0.532 | 0.732 | 0.464 | 0.015 | 0.644 | 0.152 |
| EPA_screen | 0.0608 | 0.0811 | 0.104 | 0.162 | 0.0587 | -0.0524 | 0.38 | 0.241 | 0.134 | 0.0188 | 0.396 | 0.449 |
| CDCA | 0.15 | 0.167 | 0.0118 | 0.156 | 0.171 | 0.0169 | 0.0297 | 0.0155 | 0.865 | 0.0235 | 0.0132 | 0.808 |
| TLCA | -0.185 | -0.199 | -0.0935 | -0.155 | -0.182 | -0.011 | 0.0071 | 0.0037 | 0.177 | 0.0245 | 0.0083 | 0.874 |
| ALA_screen | 0.0374 | 0.0502 | 0.0265 | 0.14 | 0.023 | -0.0389 | 0.589 | 0.468 | 0.702 | 0.0416 | 0.74 | 0.574 |
| POEA_Screen | 0.0291 | 0.0418 | 0.0593 | 0.138 | 0.0046 | -0.0321 | 0.674 | 0.546 | 0.391 | 0.045 | 0.947 | 0.643 |
| TDCA/DCA | -0.167 | -0.201 | -0.0575 | -0.134 | -0.24 | -0.0286 | 0.0152 | 0.0035 | 0.407 | 0.0521 | 0.0004 | 0.68 |
| 15-HEPE | 0.105 | 0.135 | 0.12 | 0.129 | 0.0602 | 0.0441 | 0.127 | 0.0496 | 0.0821 | 0.0612 | 0.384 | 0.524 |
| GDCA/DCA | -0.164 | -0.187 | -0.0806 | -0.126 | -0.213 | -0.0212 | 0.0173 | 0.0065 | 0.245 | 0.0689 | 0.0019 | 0.76 |
| Sum(DiHOME/EpOME) | -0.0329 | -0.1 | -0.0402 | -0.122 | -0.0785 | -0.161 | 0.635 | 0.146 | 0.561 | 0.0774 | 0.256 | 0.019 |
| TDCA/CA | -0.235 | -0.183 | -0.0247 | -0.122 | -0.127 | 0.0366 | 0.0006 | 0.0079 | 0.722 | 0.0781 | 0.0666 | 0.598 |
| TXB2 | 0.0421 | 0.0478 | 0.0466 | 0.115 | 0.0474 | -0.0496 | 0.543 | 0.489 | 0.501 | 0.0946 | 0.494 | 0.474 |
| CA | 0.183 | 0.133 | -0.0069 | 0.114 | 0.116 | -0.0611 | 0.0078 | 0.0539 | 0.92 | 0.0995 | 0.0926 | 0.378 |
| NA-Gly | 0.0193 | 0.0176 | 0.0757 | 0.106 | 0.0224 | -0.052 | 0.78 | 0.799 | 0.274 | 0.124 | 0.746 | 0.452 |
| Acetaminophen | -0.0169 | -0.108 | -0.0211 | -0.106 | -0.155 | -0.0652 | 0.807 | 0.117 | 0.761 | 0.127 | 0.024 | 0.346 |
| UDCA | -0.0682 | -0.0989 | -0.0712 | -0.104 | -0.0156 | -0.0897 | 0.326 | 0.153 | 0.304 | 0.133 | 0.822 | 0.196 |
| 9-KODE | -0.0019 | 0.0137 | -0.0898 | 0.104 | 0.028 | 0.0245 | 0.978 | 0.844 | 0.194 | 0.133 | 0.685 | 0.723 |
| CRTN | -0.0121 | -0.042 | -0.0535 | -0.104 | -0.0202 | 0.0088 | 0.861 | 0.545 | 0.44 | 0.134 | 0.772 | 0.899 |
| 14(15)-EpETrE | 0.0173 | 0.0598 | 0.074 | 0.103 | 0.0953 | 0.0346 | 0.803 | 0.387 | 0.285 | 0.135 | 0.168 | 0.617 |
| GDCA/CA | -0.232 | -0.15 | 0.0027 | -0.103 | -0.0727 | 0.0682 | 0.0007 | 0.0294 | 0.969 | 0.138 | 0.295 | 0.325 |
| DCA | 0.0623 | 0.127 | 0.036 | 0.0988 | 0.216 | 0.0578 | 0.369 | 0.0655 | 0.604 | 0.154 | 0.0017 | 0.405 |
| TCA/CA | -0.176 | -0.151 | -0.0228 | -0.0962 | -0.134 | 0.0306 | 0.0105 | 0.0286 | 0.743 | 0.165 | 0.0528 | 0.66 |
| 14_15-DiHETrE/14(15)-EpETrE | -0.0186 | -0.0638 | -0.0961 | -0.0933 | -0.0864 | -0.054 | 0.788 | 0.356 | 0.165 | 0.177 | 0.211 | 0.435 |
| GCA/CA | -0.184 | -0.131 | -0.0224 | -0.0922 | -0.0832 | 0.0719 | 0.0076 | 0.0577 | 0.747 | 0.183 | 0.23 | 0.3 |
| 12,13-DiHOME/EpOME 2 | -0.0207 | -0.0718 | 0.0041 | -0.0906 | -0.0438 | -0.145 | 0.765 | 0.299 | 0.953 | 0.19 | 0.527 | 0.0355 |
| 12_13-DiHOME/12(13)-EpOME | -0.0207 | -0.0718 | 0.0041 | -0.0906 | -0.0438 | -0.145 | 0.765 | 0.299 | 0.953 | 0.19 | 0.527 | 0.0355 |
| 9_10-e-DiHO | 0.0177 | 0.0337 | -0.028 | 0.0904 | 0.0347 | 0.0123 | 0.798 | 0.626 | 0.686 | 0.191 | 0.616 | 0.859 |
| TDCA/TLCA | 0.0339 | 0.0706 | 0.0494 | 0.0906 | 0.0813 | 0.023 | 0.625 | 0.308 | 0.476 | 0.191 | 0.241 | 0.74 |
| Sum(DiHETrE/EpETrE) | -0.0107 | -0.0485 | -0.0749 | -0.09 | -0.0802 | -0.0345 | 0.877 | 0.483 | 0.279 | 0.193 | 0.246 | 0.618 |
| w-MCA/UDCA | 0.143 | 0.155 | 0.12 | 0.0883 | 0.102 | 0.0425 | 0.0381 | 0.0247 | 0.0835 | 0.203 | 0.14 | 0.541 |
| F2-IsoP | 0.0326 | 0.0507 | 0.0847 | 0.0875 | -0.0021 | 0.0123 | 0.638 | 0.464 | 0.22 | 0.206 | 0.976 | 0.859 |
| CRCTN | -0.0139 | -0.0394 | -0.0433 | -0.087 | 0.0558 | -0.0635 | 0.841 | 0.57 | 0.533 | 0.209 | 0.421 | 0.36 |
| DGLEA | -0.0527 | -0.008 | -0.0546 | 0.0865 | 0.0332 | 0.0241 | 0.446 | 0.908 | 0.43 | 0.211 | 0.631 | 0.728 |
| 1-OG | 0.0257 | -0.0046 | -0.0121 | 0.085 | 0.0092 | -0.11 | 0.711 | 0.947 | 0.861 | 0.219 | 0.894 | 0.113 |
| EPEA_Screen | 0.0248 | 0.0288 | 0.0769 | 0.0843 | 0.018 | -0.0718 | 0.72 | 0.677 | 0.266 | 0.223 | 0.795 | 0.299 |
| 9(10)-EpOME | 0.0135 | 0.0413 | 0.0401 | 0.0842 | -0.0239 | 0.0827 | 0.846 | 0.551 | 0.562 | 0.223 | 0.73 | 0.232 |
| 17-HDoHE | -0.0414 | -0.0544 | 0.0464 | -0.082 | -0.0638 | -0.0113 | 0.55 | 0.432 | 0.503 | 0.236 | 0.356 | 0.87 |
| PGE2 | 0.027 | 0.0119 | -0.0633 | 0.0813 | 0.0336 | -0.0221 | 0.697 | 0.864 | 0.36 | 0.239 | 0.628 | 0.749 |
| TCDCA | -0.0372 | -0.0872 | -0.0625 | -0.0806 | -0.136 | -0.0177 | 0.592 | 0.208 | 0.367 | 0.245 | 0.0494 | 0.799 |
| 12(13)-EpOME | 0.007 | 0.0616 | 0.0173 | 0.0798 | 0.0258 | 0.109 | 0.919 | 0.373 | 0.803 | 0.248 | 0.71 | 0.114 |
| 11_12-DiHETrE | -0.0004 | 0.041 | 0.019 | 0.0797 | 0.0402 | 0.0376 | 0.996 | 0.554 | 0.784 | 0.249 | 0.561 | 0.588 |
| Sum(DiHODE/EpODE) | -0.0174 | -0.0205 | 0.0731 | -0.0784 | -0.0047 | -0.0816 | 0.801 | 0.767 | 0.291 | 0.257 | 0.945 | 0.238 |
| 15_16-DiHODE/15(16)-EpODE | -0.0142 | -0.0252 | 0.0625 | -0.0775 | -0.0018 | -0.0982 | 0.837 | 0.716 | 0.367 | 0.262 | 0.979 | 0.155 |

Continuation of the Supplemental Table S3

| Meatbolite | Episodic memory | Global cognition | Perceptual orientation | perceptual speed | semantic memory | working memmory | Episodic memory | Global cognition | Perceptual orientation | perceptual speed | semantic memory | working memmory |
|---|---|---|---|---|---|---|---|---|---|---|---|---|
| 9-HETE | -0.0521 | -0.0499 | -0.081 | 0.0739 | -0.0682 | -0.0311 | 0.452 | 0.471 | 0.241 | 0.285 | 0.324 | 0.653 |
| PGF2a | 0.0578 | 0.0448 | 0.072 | 0.0737 | -0.0062 | -0.0607 | 0.404 | 0.517 | 0.298 | 0.287 | 0.928 | 0.381 |
| 5_6-DiHETrE | 0.0413 | 0.0777 | 0.0843 | 0.0723 | 0.0372 | 0.0726 | 0.551 | 0.261 | 0.223 | 0.296 | 0.591 | 0.294 |
| GLCA | -0.0841 | -0.0401 | -0.0481 | -0.0723 | 0.0484 | 0.033 | 0.225 | 0.563 | 0.488 | 0.297 | 0.486 | 0.635 |
| 9,10-DiHOME/EpOME 2 | 0.016 | -0.0323 | -0.0914 | -0.0698 | 0.0129 | -0.0946 | 0.817 | 0.641 | 0.186 | 0.313 | 0.852 | 0.171 |
| 9_10-DiHOME/9(10)-EpOME | 0.016 | -0.0323 | -0.0914 | -0.0698 | 0.0129 | -0.0946 | 0.817 | 0.641 | 0.186 | 0.313 | 0.852 | 0.171 |
| CRTL | -0.0135 | -0.0428 | -0.0579 | -0.0692 | 0.0413 | -0.0482 | 0.846 | 0.537 | 0.404 | 0.318 | 0.552 | 0.487 |
| GUDCA/UDCA | 0.129 | 0.135 | 0.015 | 0.0684 | 0.0848 | 0.109 | 0.0624 | 0.0517 | 0.829 | 0.324 | 0.221 | 0.114 |
| 9-HODE | 0.0325 | 0.029 | -0.0324 | 0.0682 | -0.029 | 0.04 | 0.638 | 0.675 | 0.64 | 0.324 | 0.676 | 0.564 |
| PGE3 | 0.0174 | 0.0137 | -0.015 | -0.0633 | -0.0068 | 0.0247 | 0.802 | 0.843 | 0.829 | 0.36 | 0.922 | 0.722 |
| 5-HETE | -0.0683 | -0.0555 | -0.0062 | 0.0626 | -0.0506 | -0.0169 | 0.324 | 0.423 | 0.928 | 0.365 | 0.464 | 0.807 |
| 9_10-DiHODE | -0.0431 | -0.0209 | -0.0361 | -0.0626 | -0.0488 | 0.0426 | 0.534 | 0.762 | 0.602 | 0.366 | 0.481 | 0.539 |
| (GDCA+GLCA)/(TDCA+TLCA) | 0.191 | 0.187 | 0.0197 | 0.0626 | 0.158 | 0.0768 | 0.0055 | 0.0067 | 0.777 | 0.367 | 0.0223 | 0.268 |
| w-MCA | 0.103 | 0.115 | 0.0483 | 0.0598 | 0.153 | 0.0222 | 0.138 | 0.0961 | 0.487 | 0.389 | 0.0268 | 0.749 |
| 15_16-DiHODE | -0.0331 | -0.0066 | 0.0059 | -0.0592 | -0.0254 | 0.0286 | 0.633 | 0.924 | 0.932 | 0.392 | 0.714 | 0.68 |
| GDCA/GLCA | -0.0117 | 0.0103 | 0.0219 | 0.0588 | -0.0174 | 0.049 | 0.867 | 0.882 | 0.753 | 0.397 | 0.802 | 0.48 |
| DHEAS | -0.0161 | 0.0029 | 0.0012 | 0.0578 | 0.0563 | -0.013 | 0.817 | 0.966 | 0.987 | 0.405 | 0.417 | 0.852 |
| 14_15-DiHETrE | 0.0216 | 0.0371 | -0.0688 | 0.0575 | 0.0585 | 0.0254 | 0.755 | 0.593 | 0.32 | 0.406 | 0.398 | 0.714 |
| 1-LG | 0.0403 | -0.01 | -0.0523 | 0.0569 | -0.0026 | -0.107 | 0.56 | 0.885 | 0.45 | 0.411 | 0.97 | 0.122 |
| TDCA | -0.141 | -0.117 | -0.0176 | -0.0559 | -0.0876 | 0.0031 | 0.0417 | 0.0917 | 0.8 | 0.421 | 0.206 | 0.964 |
| NO-Gly | 0.009 | -0.0006 | 0.0178 | 0.0552 | -0.0069 | -0.0603 | 0.897 | 0.993 | 0.797 | 0.425 | 0.92 | 0.384 |
| 5-HEPE | 0.0415 | 0.0169 | 0.12 | 0.0538 | -0.0376 | -0.088 | 0.548 | 0.807 | 0.0816 | 0.437 | 0.587 | 0.203 |
| DHEA | 0.0513 | 0.0311 | 0.055 | 0.0521 | -0.0234 | -0.025 | 0.458 | 0.654 | 0.427 | 0.452 | 0.736 | 0.718 |
| 4-HDoHE | 0.0461 | 0.0549 | 0.0219 | 0.0513 | -0.0066 | 0.0498 | 0.506 | 0.427 | 0.752 | 0.459 | 0.924 | 0.472 |
| 9-KODE/9-HODE | -0.0158 | 0.019 | -0.0283 | 0.0512 | 0.0862 | 0.0215 | 0.819 | 0.784 | 0.683 | 0.46 | 0.213 | 0.756 |
| 12-HETE | 0.0084 | -0.0311 | 0.0101 | -0.0509 | -0.0617 | -0.0073 | 0.903 | 0.653 | 0.884 | 0.462 | 0.373 | 0.916 |
| 2-AG | 1 | -0.0497 | -0.0275 | 0.0501 | -0.0132 | -0.146 | 0.999 | 0.473 | 0.691 | 0.469 | 0.849 | 0.0341 |
| AEA | -0.073 | -0.0552 | -0.001 | 0.0458 | -0.0635 | -0.0237 | 0.292 | 0.425 | 0.989 | 0.509 | 0.359 | 0.732 |
| PGD2 | 0.0564 | 0.0883 | 0.0343 | 0.0457 | 0.141 | 0.0286 | 0.415 | 0.201 | 0.621 | 0.509 | 0.0403 | 0.679 |
| 13-HODE | 0.0201 | 0.0127 | 0.0122 | 0.0449 | -0.0716 | 0.0245 | 0.771 | 0.854 | 0.86 | 0.517 | 0.301 | 0.724 |
| aLEA | -0.0091 | -0.0154 | 0.0147 | -0.0435 | 0.0152 | -0.0239 | 0.896 | 0.824 | 0.832 | 0.53 | 0.826 | 0.73 |
| DCA/CA | -0.166 | -0.062 | 0.0289 | -0.0423 | 0.0331 | 0.104 | 0.0159 | 0.371 | 0.677 | 0.542 | 0.634 | 0.133 |
| 2-OG | 0.0508 | 0.011 | 0.0029 | 0.0413 | 0.0059 | -0.073 | 0.463 | 0.874 | 0.967 | 0.551 | 0.932 | 0.291 |
| Aspirin | 0.0288 | 0.0244 | 0.0393 | 0.0401 | -0.0502 | 0.0505 | 0.678 | 0.725 | 0.57 | 0.562 | 0.468 | 0.465 |
| GUDCA | 0.048 | 0.0323 | -0.0978 | -0.0396 | 0.0929 | 0.032 | 0.489 | 0.642 | 0.158 | 0.568 | 0.18 | 0.645 |
| 12_13-DiHODE/12(13)-EpODE | 0.126 | 0.0828 | 0.124 | 0.0378 | -0.0749 | 0.0565 | 0.0676 | 0.231 | 0.0734 | 0.585 | 0.279 | 0.415 |
| DEA | -0.0245 | -0.0662 | -0.0434 | 0.0374 | -0.0492 | -0.114 | 0.723 | 0.338 | 0.531 | 0.589 | 0.477 | 0.0983 |
| 2-LG | 0.0243 | -0.0292 | -0.054 | 0.0371 | -0.0131 | -0.113 | 0.726 | 0.673 | 0.435 | 0.592 | 0.85 | 0.101 |
| 12-HEPE | 0.0131 | -0.0191 | 0.0381 | -0.0354 | -0.0852 | -0.0022 | 0.85 | 0.783 | 0.582 | 0.61 | 0.218 | 0.974 |
| 13-HOTE | -0.0107 | 0.0237 | 0.037 | 0.035 | -0.008 | 0.0607 | 0.878 | 0.732 | 0.593 | 0.613 | 0.908 | 0.38 |
| 12-HEPE/12-HETE | 0.0134 | 0.0415 | 0.0624 | 0.0348 | -0.0322 | 0.0472 | 0.847 | 0.549 | 0.367 | 0.615 | 0.642 | 0.495 |
| TCA | -0.044 | -0.0559 | -0.0218 | -0.0335 | -0.0834 | 0.0201 | 0.526 | 0.421 | 0.753 | 0.629 | 0.229 | 0.772 |
| 9_10-DiHOME | -0.0217 | -0.0319 | -0.0424 | -0.0331 | -0.0748 | 0.02 | 0.754 | 0.645 | 0.541 | 0.633 | 0.28 | 0.773 |
| GCDCA/GLCA | 0.0776 | 0.0253 | 0.0349 | 0.0332 | -0.0678 | -0.0255 | 0.263 | 0.715 | 0.615 | 0.633 | 0.328 | 0.713 |
| 1-AG | 0.0041 | -0.0517 | -0.0413 | 0.0326 | -0.0218 | -0.124 | 0.953 | 0.455 | 0.551 | 0.638 | 0.752 | 0.0716 |
| GCA/GDCA | 0.0667 | 0.0269 | -0.0082 | 0.0324 | -0.0359 | -0.002 | 0.336 | 0.699 | 0.906 | 0.64 | 0.605 | 0.977 |
| 11-Deoxy-CTRL | -0.0059 | -0.0214 | -0.0017 | -0.0324 | -0.0173 | -0.0426 | 0.932 | 0.758 | 0.981 | 0.641 | 0.803 | 0.54 |
| 15(16)-EpODE | 0.0075 | 0.0502 | -0.0351 | 0.0316 | 0.0251 | 0.112 | 0.914 | 0.468 | 0.612 | 0.648 | 0.717 | 0.106 |
| 11-HETE | 0.0394 | 0.0116 | 0.0107 | 0.0299 | 0.0025 | -0.0414 | 0.569 | 0.868 | 0.877 | 0.666 | 0.971 | 0.55 |
| OEA | 0.0183 | 0.0049 | 0.025 | 0.0298 | -0.0137 | -0.0745 | 0.792 | 0.943 | 0.718 | 0.667 | 0.844 | 0.282 |
| 9_10-DiHODE/9(10)-EpODE | -0.0331 | 0.009 | 0.0222 | -0.0287 | 0.038 | 0.0238 | 0.633 | 0.897 | 0.748 | 0.679 | 0.583 | 0.731 |
| 20-HETE | -0.0562 | -0.0429 | 0.014 | 0.027 | 0.0779 | -0.151 | 0.417 | 0.535 | 0.839 | 0.696 | 0.26 | 0.0286 |
| 9-HOTE | 0.0372 | 0.0523 | -0.0075 | 0.0262 | 0.0195 | 0.0488 | 0.591 | 0.45 | 0.914 | 0.705 | 0.778 | 0.481 |
| 9_12_13-TriHOME | 0.0468 | 0.0081 | -0.0186 | 0.0259 | -0.0345 | -0.0882 | 0.499 | 0.907 | 0.788 | 0.709 | 0.618 | 0.202 |
| 14_15-DiHETE | -0.0172 | -0.0202 | 0.0659 | -0.0254 | -0.0734 | 0.0039 | 0.804 | 0.771 | 0.341 | 0.714 | 0.289 | 0.955 |
| Sum(HDoHEs) | -0.0009 | -0.0323 | 0.0501 | -0.024 | -0.0889 | -0.0181 | 0.99 | 0.641 | 0.47 | 0.729 | 0.199 | 0.794 |
| 14,15/11,12-DiHETrE | 0.059 | -0.0012 | -0.12 | -0.0212 | 0.0264 | -0.0653 | 0.394 | 0.986 | 0.0823 | 0.76 | 0.704 | 0.345 |
| 14-HDoHE | 0.0022 | -0.0272 | 0.0567 | -0.0196 | -0.0874 | -0.0133 | 0.974 | 0.695 | 0.413 | 0.778 | 0.206 | 0.848 |
| 17OH-PROG | 0.0538 | 0.0356 | -0.0813 | 0.0196 | 0.0248 | 0.0293 | 0.438 | 0.609 | 0.241 | 0.778 | 0.721 | 0.673 |
| 15-HETE | 0.0014 | -0.0171 | 0.0216 | 0.0193 | -0.0493 | -0.0142 | 0.984 | 0.805 | 0.755 | 0.78 | 0.476 | 0.837 |

Continuation of the Supplemental Table S3

| Meatbolite | Episodic memory | Global cognition | Perceptual orientation | perceptual speed | semantic memory | working memmory | Episodic memory | Global cognition | Perceptual orientation | perceptual speed | semantic memory | working memmory |
|---|---|---|---|---|---|---|---|---|---|---|---|---|
| 8_9-DiHETrE | -0.0538 | -0.0524 | -0.06 | -0.0172 | -0.0379 | -0.0236 | 0.437 | 0.449 | 0.386 | 0.804 | 0.584 | 0.734 |
| 12_13-DiHOME | -0.0171 | -0.0087 | 0.0116 | -0.0168 | -0.0467 | 0.0258 | 0.805 | 0.9 | 0.867 | 0.808 | 0.5 | 0.71 |
| 17_18-DiHETE | 0.0047 | 0.0173 | 0.0054 | -0.0168 | -0.0351 | 0.047 | 0.946 | 0.802 | 0.938 | 0.809 | 0.612 | 0.497 |
| Sum(DiHETEs) | -0.0009 | 0.011 | 0.0167 | -0.0147 | -0.0376 | 0.0397 | 0.989 | 0.874 | 0.81 | 0.832 | 0.587 | 0.566 |
| Sum_n3_Diols | -0.0219 | -0.003 | 0.013 | -0.0146 | -0.0436 | 0.0381 | 0.752 | 0.966 | 0.851 | 0.833 | 0.529 | 0.582 |
| 17_18_DiHETE+19_20_DiHDoPe | -0.0158 | 0.0038 | 0.0066 | -0.0144 | -0.0401 | 0.042 | 0.82 | 0.956 | 0.924 | 0.835 | 0.563 | 0.544 |
| Average n3 diols | -0.0158 | 0.0038 | 0.0066 | -0.0144 | -0.0401 | 0.042 | 0.82 | 0.956 | 0.924 | 0.835 | 0.563 | 0.544 |
| Tes/Prog | -0.0038 | 0.0094 | 0.0743 | -0.0144 | -0.0562 | 0.0279 | 0.956 | 0.893 | 0.284 | 0.836 | 0.418 | 0.687 |
| GCDCA | 0.0194 | 0.0049 | -0.0334 | -0.0143 | 0.0093 | 0.0053 | 0.779 | 0.944 | 0.631 | 0.837 | 0.893 | 0.939 |
| LTB4 | -0.0993 | -0.0704 | -0.0415 | 0.0113 | -0.0953 | 0.0211 | 0.151 | 0.309 | 0.549 | 0.87 | 0.168 | 0.76 |
| GDCA | -0.102 | -0.0403 | -0.0131 | -0.0098 | 0.0379 | 0.0404 | 0.142 | 0.561 | 0.851 | 0.888 | 0.585 | 0.561 |
| CA/CDCA | 0.074 | -0.0165 | -0.0216 | -0.008 | -0.0585 | -0.111 | 0.286 | 0.813 | 0.756 | 0.908 | 0.399 | 0.108 |
| GCA | -0.0179 | -0.0034 | -0.0175 | 0.0069 | 0.014 | 0.0323 | 0.797 | 0.962 | 0.801 | 0.921 | 0.84 | 0.642 |
| 5_15-DiHETE | -0.039 | 0.0089 | 0.123 | 0.0065 | 0.0548 | -0.0145 | 0.573 | 0.898 | 0.0739 | 0.926 | 0.428 | 0.834 |
| LEA | -0.0229 | -0.0412 | -0.0721 | -0.006 | -0.0123 | -0.0398 | 0.741 | 0.552 | 0.297 | 0.931 | 0.859 | 0.565 |
| TEST | 0.0446 | 0.0492 | -0.0002 | 0.0059 | -0.0149 | 0.0606 | 0.52 | 0.478 | 0.998 | 0.933 | 0.83 | 0.382 |
| 9(10)-EpODE | -0.044 | -0.0678 | -0.0546 | 0.0055 | -0.0866 | -0.0537 | 0.526 | 0.327 | 0.43 | 0.937 | 0.21 | 0.438 |
| 12_13-DiHODE | 0.0753 | 0.0922 | 0.0618 | 0.0036 | 0.0265 | 0.101 | 0.276 | 0.182 | 0.372 | 0.958 | 0.702 | 0.143 |
| PGE2/PGD2 | 0.0942 | 0.114 | 0.107 | 0.0033 | 0.087 | 0.0596 | 0.173 | 0.0999 | 0.123 | 0.962 | 0.208 | 0.389 |
| PGF2a-1G | -0.0392 | -0.0543 | -0.128 | 0.0033 | -0.0196 | -0.0852 | 0.571 | 0.433 | 0.0646 | 0.962 | 0.777 | 0.218 |
| 12(13)-EpODE | -0.0343 | 0.0317 | -0.0453 | -0.0025 | 0.112 | 0.0846 | 0.62 | 0.647 | 0.513 | 0.971 | 0.105 | 0.221 |
| Ibuprofen | -0.103 | -0.0993 | -0.038 | 0.0023 | -0.0313 | -0.105 | 0.161 | 0.177 | 0.606 | 0.975 | 0.671 | 0.154 |
| 19_20-DiHDoPE | -0.059 | -0.0276 | 0.0222 | -0.0017 | -0.0507 | 0.0127 | 0.394 | 0.69 | 0.749 | 0.98 | 0.464 | 0.854 |
