## Supplemental Table S4 for "Serum metabolomic biomarkers of perceptual speed in cognitively normal and mildly impaired subjects with fasting state stratification"

**Supplemental Table S4.** Stepwise linear model predictive cognitive domains, stratified by the fasting state. Stepwise analysis was performed with the maximal validation r² as the model stopping criteria, or if an additional step increased the BIC. Model stopping point for each domain is highlighted.

### Fasted

| Cognitive domain | Step | Parameter | Sig Prob | Seq SS | RSquare | AICc | BIC | RSquare Validation |
|---|---|---|---|---|---|---|---|---|
| **Episodic memory** | 1 | **Johnson Su[LTB4] 2** | **0.0083** | 2.34 | 0.174 | 68.4 | 72.7 | -0.12 |
| | 2 | **Johnson Su[T-a-MCA]** | **0.0214** | 1.54 | 0.288 | 65.1 | 70.6 | -0.0568 |
| | 3 | Johnson Sb[GUDCA/UDCA] | 0.131 | 0.613 | 0.334 | 65.2 | 71.7 | 0.127 |
| | 4 | Johnson Su[19_20-DiHDoPE] 2 | 0.299 | 0.285 | 0.355 | 66.7 | 74.1 | 0.14 |
| **Perceptual orientation** | 1 | **Johnson Sb[TDCA/CA]** | **0.0106** | 2.34 | 0.164 | 71.2 | 75.5 | -0.0762 |
| | 2 | **Johnson Sb[GUDCA/UDCA]** | **0.0729** | 1.03 | 0.236 | 70.1 | 75.6 | 0.0806 |
| | 3 | **Johnson Su[PGF2a-1G] 2** | **0.036** | 1.3 | 0.328 | 67.8 | 74.3 | 0.222 |
| | 4 | **PGE2/PGD2** | **0.0572** | 0.982 | 0.397 | 66.4 | 73.8 | 0.229 |
| | 5 | Johnson Su[9-HETE] 2 | 0.156 | 0.517 | 0.433 | 67 | 75 | 0.254 |
| | 6 | Johnson Su[GDCA/DCA] | 0.186 | 0.437 | 0.463 | 68 | 76.5 | 0.249 |
| **Perceptual speed** | 1 | **Johnson Sb[12,1...DiHOME/EpOME 2]** | **0.0014** | 5 | 0.219 | 91.7 | 96.5 | -0.027 |
| | 2 | **Johnson Su[Sum_n3_Diols]** | **0.0018** | 3.81 | 0.386 | 83.6 | 89.7 | 0.0919 |
| | 3 | **Johnson Su[T-a-MCA]** | **0.0462** | 1.34 | 0.445 | 81.7 | 89 | 0.235 |
| | 4 | Johnson Su[PGF2a] | 0.38 | 0.251 | 0.456 | 83.5 | 91.9 | 0.302 |
| | 5 | Johnson Su[19_20-DiHDoPE] | 0.585 | 0.0984 | 0.46 | 86 | 95.4 | 0.34 |
| **Semantic memory** | 1 | Johnson Su[17-HDoHE] | 0.0458 | 1.36 | 0.104 | 70.8 | 75.1 | -0.315 |
| | 2 | Johnson Su[T-a-MCA] | 0.0232 | 1.6 | 0.225 | 67.6 | 73.1 | -0.332 |
| | 3 | Johnson Sb[TXB2] | 0.0638 | 0.968 | 0.298 | 66.4 | 72.9 | -0.477 |
| **Working memory** | 1 | **Johnson Su[CRTN]** | 0.127 | 0.922 | 0.062 | 77.3 | 81.6 | 0.0093 |
| | 2 | Johnson Su[15_1...E/15(16)-EpODE] | 0.206 | 0.615 | 0.103 | 78 | 83.5 | 0.0736 |
| | 3 | Johnson Sb[ALA_screen] | 0.407 | 0.264 | 0.121 | 79.9 | 86.4 | -0.0302 |
| **Global cognition** | 1 | Johnson Sb[GUDCA/UDCA] | **0.0244** | 1.23 | 0.115 | 64 | 68.7 | -0.252 |
| | 2 | **Johnson Su[T-a-MCA]** | **0.0137** | 1.32 | 0.238 | 59.8 | 65.9 | 0.127 |
| | 3 | Johnson Su[15(16)-EpODE] | 0.101 | 0.539 | 0.289 | 59.4 | 66.7 | -0.0067 |
| | 4 | Johnson Su[19_20-DiHDoPE] | 0.165 | 0.373 | 0.323 | 59.8 | 68.3 | 0.0578 |

### Non-fasted

| Cognitive domain | Step | Parameter | Sig Prob | Seq SS | RSquare | AICc | BIC | RSquare Validation |
|---|---|---|---|---|---|---|---|---|
| **Episodic memory** | 1 | **Johnson Sb[TLCA]** | **0.0003** | 3.94 | 0.144 | 139 | 146 | -0.011 |
| | 2 | Johnson Sb[4-HDoHE] | **0.0258** | 1.35 | 0.194 | 136 | 145 | -0.196 |
| | 3 | Johnson Su[12_1...E/12(13)-EpODE] | 0.153 | 0.538 | 0.213 | 136 | 148 | -0.142 |
| **Perceptual orientation** | 1 | **Johnson Sb[DCA/CA]** | **0.0573** | 1.7 | 0.0419 | 183 | 190 | -0.0227 |
| | 2 | Johnson Sb[TDCA/TLCA] | 0.0581 | 1.64 | 0.0822 | 182 | 191 | -0.1 |
| | 3 | Johnson Sb[5-HEPE] | 0.14 | 0.973 | 0.106 | 182 | 193 | -0.12 |
| **Perceptual speed** | 1 | **Johnson Su[LA_screen]** | **0.0071** | 3.98 | 0.0821 | 195 | 202 | -0.128 |
| | 2 | Johnson Sb[4-HDoHE] | **0.0343** | 2.32 | 0.13 | 192 | 202 | -0.218 |
| | 3 | Johnson Sb[TXB2] | 0.232 | 0.724 | 0.145 | 193 | 205 | -0.291 |
| **Semantic memory** | 1 | **Johnson Su[GDCA/DCA]** | **0.0002** | 3.59 | 0.153 | 125 | 132 | -0.107 |
| | 2 | Johnson Su[12(13)-EpODE] | **0.0402** | 0.978 | 0.194 | 123 | 132 | -0.0327 |
| | 3 | Johnson Sb[TLCA] | 0.104 | 0.597 | 0.22 | 122 | 134 | 0.002 |
| **Working memory** | 1 | **Johnson Sb[AA_screen]** | **0.0243** | 2.08 | 0.0583 | 170 | 177 | 0.0233 |
| | 2 | Johnson Su[(GDC...A)/(TDCA+TLCA)] | 0.0594 | 1.4 | 0.0975 | 169 | 178 | -0.0345 |
| | 3 | Johnson Sb[Tes/Prog] | 0.198 | 0.641 | 0.116 | 169 | 181 | -0.114 |
| **Global cognition** | 1 | **Johnson Sb[TLCA]** | **0.0002** | 2.36 | 0.152 | 89.2 | 96.4 | -0.104 |
| | 2 | Johnson Su[12_13-DiHODE] | **0.0475** | 0.607 | 0.191 | 87.4 | 96.7 | -0.0725 |
| | 3 | Johnson Su[GDCA/DCA] | 0.274 | 0.181 | 0.202 | 88.3 | 99.9 | -0.0481 |
